# Nvj3 regulates Dga1-mediated triacylglycerol synthesis and lipid droplet formation at ER contact sites

**DOI:** 10.64898/2026.01.05.695145

**Authors:** Daniel Adebayo, Eseiwi Obaseki, Scott Miller, Marwa Aboumourad, Zaid Saadeh, Zachary Pomicter, Lindsey Kreinbring, Mohamad Rahal, Kashvi Vasudeva, Garrett Frost, Reema Smadi, Hanaa Hariri

## Abstract

Lipid droplet (LD) biogenesis is essential for lipid homeostasis during nutrient stress, yet how lipid intermediates are spatially organized to support efficient triacylglycerol (TAG) synthesis remains unclear. Here, we identify Nvj3 as a nutrient-responsive regulator that links diacylglycerol (DAG) availability to TAG synthesis and LD formation at the endoplasmic reticulum (ER). Nvj3 is induced by glucose depletion and recruited to LD-associated ER domains. Loss of Nvj3 causes neutral lipid accumulation under steady state conditions but delays TAG synthesis under acute inducible metabolic transitions. Using controlled TAG induction systems, we show that Nvj3 is required to couple Dga1-dependent TAG synthesis to LD formation. In the absence of Nvj3, TAG accumulates but remains inefficiently packaged into LDs. Consistent with this defect, *nvj3Δ* cells exhibit altered phospholipid remodeling and mislocalization of DAG away from ER domains during starvation. Together, these findings establish Nvj3 as an organizer of lipid availability during metabolic stress and suggest that spatial control of DAG is a key determinant of LD biogenesis.

**Summary:** This study identifies Nvj3 as a spatial organizer of Dga1-dependent lipid droplet formation. Nvj3 promotes proper diacylglycerol positioning, and enables efficient triacylglycerol synthesis during metabolic stress. We propose that Nvj3 regulates lipid flux through spatial compartmentalization of diacylglycerol at membrane contact site-associated ER domains.

## Introduction

Cells respond to nutrient stress by remodeling their metabolic networks to maintain lipid homeostasis. A critical process in this adaptation involves the storage of excess free fatty acids (FFAs) in the form of triacylglycerol (TAG) within lipid droplets (LDs); dynamic organelles that form at the endoplasmic reticulum (ER) (Olzmann and Carvalho, 2019; Walther and Farese, 2012). LDs buffer excess FFAs, protecting cells from lipotoxicity while also serving as metabolically active reservoirs that can be rapidly mobilized in response to changing nutrient conditions (Obaseki et al., 2024; Petschnigg et al., 2009; Listenberger et al., 2003). TAG within LDs can be hydrolyzed to generate FFAs and diacylglycerol (DAG) for energy production and membrane biosynthesis (Grabner et al., 2021; Graef, 2018; Singh et al., 2009). Therefore, LD biogenesis and turnover are tightly coupled to coordinate lipid storage with membrane synthesis, particularly during nutrient limitation (Lysyganicz et al., 2025; Barbosa and Siniossoglou, 2016; Barbosa et al., 2015).

Several enzymes involved in neutral lipid synthesis are localized to the ER, where the availability and routing of lipid intermediates are critical for LD biogenesis. In budding yeast (*Saccharomyces cerevisiae*), DAG is primarily synthesized from phosphatidic acid (PA) by the phosphatase Pah1 (Adeyo et al., 2011). Conversion of DAG into TAG is catalyzed by two enzymes with distinct topologies and substrate requirements: Dga1, a cytosolic-facing acyl-CoA-dependent diacylglycerol acyltransferase, and Lro1, a luminal ER phospholipid:diacylglycerol acyltransferase (Dahlqvist et al., 2000; Farese and Walther, 2023; Buhman et al., 2001). These enzymes contribute to TAG synthesis in a growth-phase-dependent manner (Oelkers et al., 2002). Lro1 plays a dominant role during exponential growth, whereas Dga1 becomes the principal TAG synthase during stationary phase and nutrient stress, coinciding with its relocalization from the ER to LDs where it drives LD expansion (Jacquier et al., 2011; Markgraf et al., 2014; Wilfling et al., 2013). Although both Dga1 and Lro1 utilize DAG as a substrate, their distinct topologies imply fundamentally different modes of access to DAG pools within the ER. Yet, how DAG pools are spatially organized and maintained at ER sites where LD formation takes place remains unclear.

LDs do not form randomly within the ER but instead emerge from specialized ER subdomains demarcated by specific proteins (Joshi et al., 2021; Choudhary et al., 2020). LD biogenesis is also strongly governed by the biophysical properties of the ER membrane, and the spatial dynamics of lipid flux (Thiam et al., 2013; Elhan et al., 2025; Zoni et al., 2021; Ben M’barek et al., 2017). Extensive biophysical and *in vitro* reconstitution studies have shown that newly synthesized TAG “demixes” from the ER bilayer to form neutral lipid “lenses” once a local concentration threshold of ∼3 mol% is reached, initiating nascent LD formation (Choudhary et al., 2015; Walther et al., 2023). The rate and localization of TAG synthesis, rather than bulk TAG levels, determine whether TAG is incorporated into pre-existing LDs (Choudhary et al., 2018; Thiam et al., 2013; Choudhary et al., 2015). Consistent with this, recent work has shown that lipid transfer proteins can directly gate access of DAG to TAG synthases. In mammalian cells, ATG2-mediated DAG transfer was shown to recruit DGAT2 (Dga1 in yeast) and promote LD growth, demonstrating that spatial delivery of DAG can be rate-limiting for productive TAG synthesis (Elhan et al., 2025). These findings suggest that regulated positioning of lipid intermediates, rather than enzyme abundance alone, may represent a conserved principle governing LD biogenesis.

Membrane lipid composition further modulates LD biogenesis by shaping local curvature and surface tension of the ER membrane (Ben M’barek et al., 2017; Thiam et al., 2013). Lipids exhibiting a negative intrinsic curvature such as DAG and PE (phosphatidylethanolamine), promote LD nucleation and stabilize the highly curved ER-LD interface facilitating TAG lens formation (Choudhary et al., 2018; Chorlay and Thiam, 2018). Conversely, the incorporation of saturated and short-chain acyl chains into membrane lipids tends to oppose TAG lens formation, promoting neutral lipid accumulation with the ER bilayer (Zoni et al., 2021). In contrast, enrichment of positively curved lipids, specifically lysophospholipids (LPA and LPC) stabilize emerged LD state and promote their budding (Choudhary et al., 2018; Chorlay and Thiam, 2018). These findings collectively demonstrate that efficient LD formation requires not only TAG synthesis, but precise spatial coordination of lipid intermediates and membrane composition at specialized ER subdomains.

LDs do not function in isolation but are embedded within dynamic networks of membrane contact sites (MCS) that coordinate lipid synthesis, storage, and exchange (Renne and Hariri, 2021; Barbosa and Siniossoglou, 2017; Schuldiner and Bohnert, 2017). In yeast, one such site is the nuclear ER-vacuole junction (NVJ); an ER-vacuole contact site that expands during nutrient limitation and has emerged as a central hub for stress signaling and lipid metabolism (Pan et al., 2000; Hariri et al., 2018; Kvam and Goldfarb, 2006). LDs frequently cluster at the NVJ during starvation forming tri-organelle contacts (LD-ER-Vacuole) that support both neutral lipid synthesis and lipophagic turnover (Hariri et al., 2019; Álvarez-Guerra et al., 2024; Diep et al., 2024; Barbosa and Siniossoglou, 2016; Murphy et al., 2009).

The NVJ is classically tethered by the interaction between the ER protein Nvj1 and the vacuolar protein Vac8, but additional components contribute to its metabolic function (Pan et al., 2000; Kvam and Goldfarb, 2006). Among these are the PX-domain-containing protein Mdm1 and its paralogue Nvj3 which belong to the conserved sorting nexin–regulator of G protein signaling (SNX-RGS) protein sub-family (Henne et al., 2015). SNX-RGS proteins are molecular tethers localized to multiple inter-organelle contact sites that exhibit roles in cellular metabolism (Hariri and Henne, 2022; Datta et al., 2019; Ugrankar et al., 2019). Mdm1 and Nvj3 localize to the NVJ independently of Nvj1 (Henne et al., 2015). Mdm1 is an ER-anchored transmembrane protein that binds vacuolar PI3P and promotes LD biogenesis during stress by locally concentrating FA activation through recruitment of the FA-CoA synthase, Faa1 (Hariri et al., 2018, 2019; Datta et al., 2019, 2020; Hariri and Henne, 2022). Recent structural modeling has suggested that Mdm1 contains a hydrophobic tunnel reminiscent of lipid transfer proteins, raising the possibility that it participates directly in channeling lipid intermediates at the ER-vacuole interfaces (Paul et al., 2022; Wong et al., 2019; Castro et al., 2022). Intriguingly, Nvj3 shares a similar predicted architecture, including a central hydrophobic cavity, suggesting a potential capacity for lipid transfer (Paul et al., 2022; Castro et al., 2022). However, the role of Nvj3 in lipid metabolism, or how it functionally interacts with Mdm1 and the TAG synthesis pathways, has remained unknown.

In this study, we investigate the role of Nvj3 in lipid remodeling and stress adaptation. We identify Nvj3 as a nutrient-responsive organizer of TAG synthesis and LD formation during glucose starvation. Loss of Nvj3 leads to aberrant DAG accumulation in the vacuole, altered phospholipid profile, and delayed TAG synthesis during inducible metabolic transitions. These defects preferentially impact the Dga1-dependent pathway, indicating a specific requirement for Nvj3 in coordinating spatial availability of DAG within ER-membranes to support TAG synthesis. Our findings support a model in which Nvj3 regulates the spatial accessibility of lipid intermediates at ER contact sites, enabling efficient coupling of TAG synthesis to LD biogenesis during metabolic stress. Together, these observations raise the possibility that stress-induced regulators at ER contact sites control LD formation by governing access of TAG synthases to spatially restricted DAG pools.

## Results

### Nvj3 is induced by glucose depletion and relocalizes to LDs at the NVJ

To investigate the role of Nvj3 in adaptive lipid remodeling during nutrient stress, we first examined its expression and localization in response to glucose depletion. Compared to log phase cultures, we found that Nvj3 mRNA levels were upregulated ∼5-fold both in stationary phase (SP; gradual starvation) and in acute glucose starvation (AGS), consistent with a potential role in metabolic adaptation (Figure 1a).

**Figure 1.**
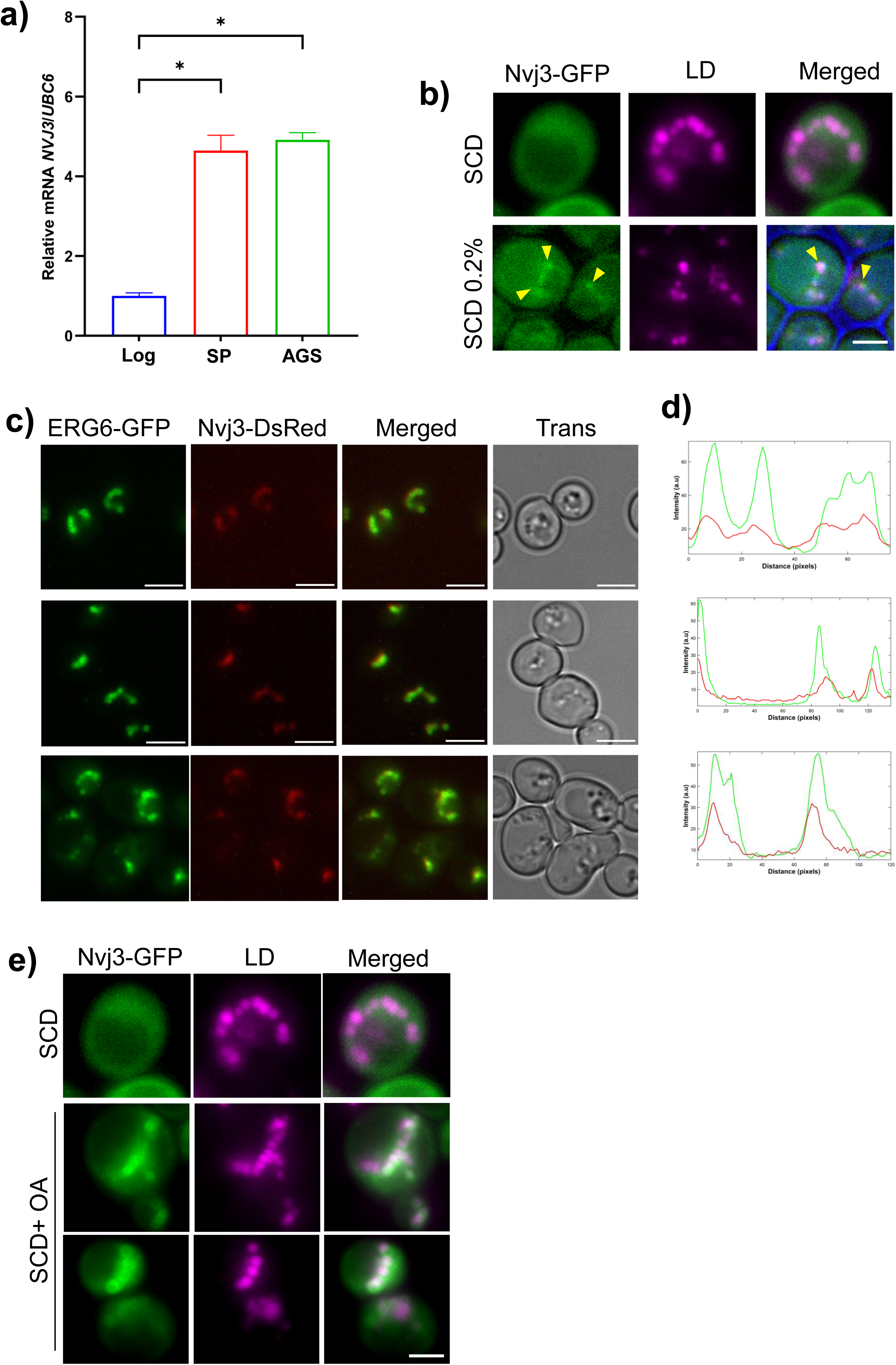
Nvj3 is a starvation-induced protein that localizes to LDs at the NVJ. A. Quantitative qPCR analysis of *NVJ3* mRNA levels in log phase, stationary phase (SP, 72h), and acute glucose starvation (AGS; 0.001% glucose, 6h). Expression is normalized to *UBC6* and plotted relative to log phase. Bars represent mean ± SEM from n = 4 biological replicates; * *P < 0.05* by unpaired t test. B. Fluorescence imaging of endogenously tagged Nvj3–GFP (green) and LDs (magenta, MDH) in cells grown in SCD media (2% or 0.2% dextrose). Arrowheads mark Nvj3–GFP puncta colocalizing with LDs at the NVJ. Scale bar, 1 µm. C. Fluorescence imaging of endogenously-tagged Nvj3–DsRed with the LD marker Erg6–GFP in SP (48h) yeast showing colocalization. Scale bar, 3 µm. D. Line scans of fluorescence intensity from representative cells in (c) showing coincident Erg6–GFP (green) and Nvj3–DsRed (red) signals. E. Fluorescence imaging of overexpressed Nvj3-GFP showing enhanced LD association under oleate-induced conditions. Nvj3-GFP (green) and LDs (magenta, MDH) were visualized in cells grown in SCD or SCD + 0.2% oleic acid (OA). Scale bar, 1 µm.

Fluorescent live-cell imaging further revealed that glucose limitation (0.2% glucose) elicited a pronounced redistribution of endogenous Nvj3-GFP from diffuse cytosolic pattern into discrete foci at the NVJ (Figure 1b), a subdomain previously identified as hotspots for LD biogenesis (Hariri et al., 2018). Under these conditions, Nvj3-GFP colocalized with LDs marked by MDH staining, indicating that starvation promotes colocalization of Nvj3 with LDs forming at NVJ-associated ER domains (Figure 1b).

To validate LD association independent of the NVJ context, we assessed Nvj3 colocalization with Erg6, a bona fide LD marker, and observed robust overlap between Erg6-GFP and endogenous Nvj3-DsRed in SP (Figure 1c, d). Furthermore, oleate (OA) supplementation, which stimulates TAG synthesis and LD expansion, enhanced Nvj3 recruitment to LDs (Figure 1e), supporting a model in which Nvj3 is dynamically targeted to LD-containing ER domains in response to lipid metabolic state of the cell.

Because prior work established that Nvj3 targeting to the NVJ requires the tethering protein Mdm1 (Henne et al., 2015), we asked whether Nvj3-LD association also depends on Mdm1. In *mdm1Δ* cells stimulated with OA, Nvj3 retained clear colocalization with LDs, demonstrating that LD association is an intrinsic property of Nvj3 and could be mechanistically distinct from its Mdm1-dependent NVJ targeting (Figure S1a).

Together, these observations show that Nvj3 is a starvation-induced protein that can be recruited to LDs in response to metabolic cues. We therefore focused subsequent experiments on determining how Nvj3 impacts LD metabolism and lipid remodeling during glucose starvation.

### Nvj3 deletion causes accumulation of LDs and neutral lipids under nutrient stress

To determine whether starvation-induced Nvj3 relocalization reflects a functional role in lipid storage, we examined LD abundance in *nvj3*Δ cells under gradual starvation (SP) and AGS (0.001% glucose). In both conditions, BODIPY staining revealed a marked increase in LD numbers relative to wild-type controls (Figure 2a, b), which was corroborated by quantitative imaging of Erg6-positive LDs (Figure S1b, c). Importantly, overexpression of Nvj3 suppressed LD accumulation, confirming that the altered LD levels resulted from the loss of Nvj3 (Figure 2c).

**Figure 2:**
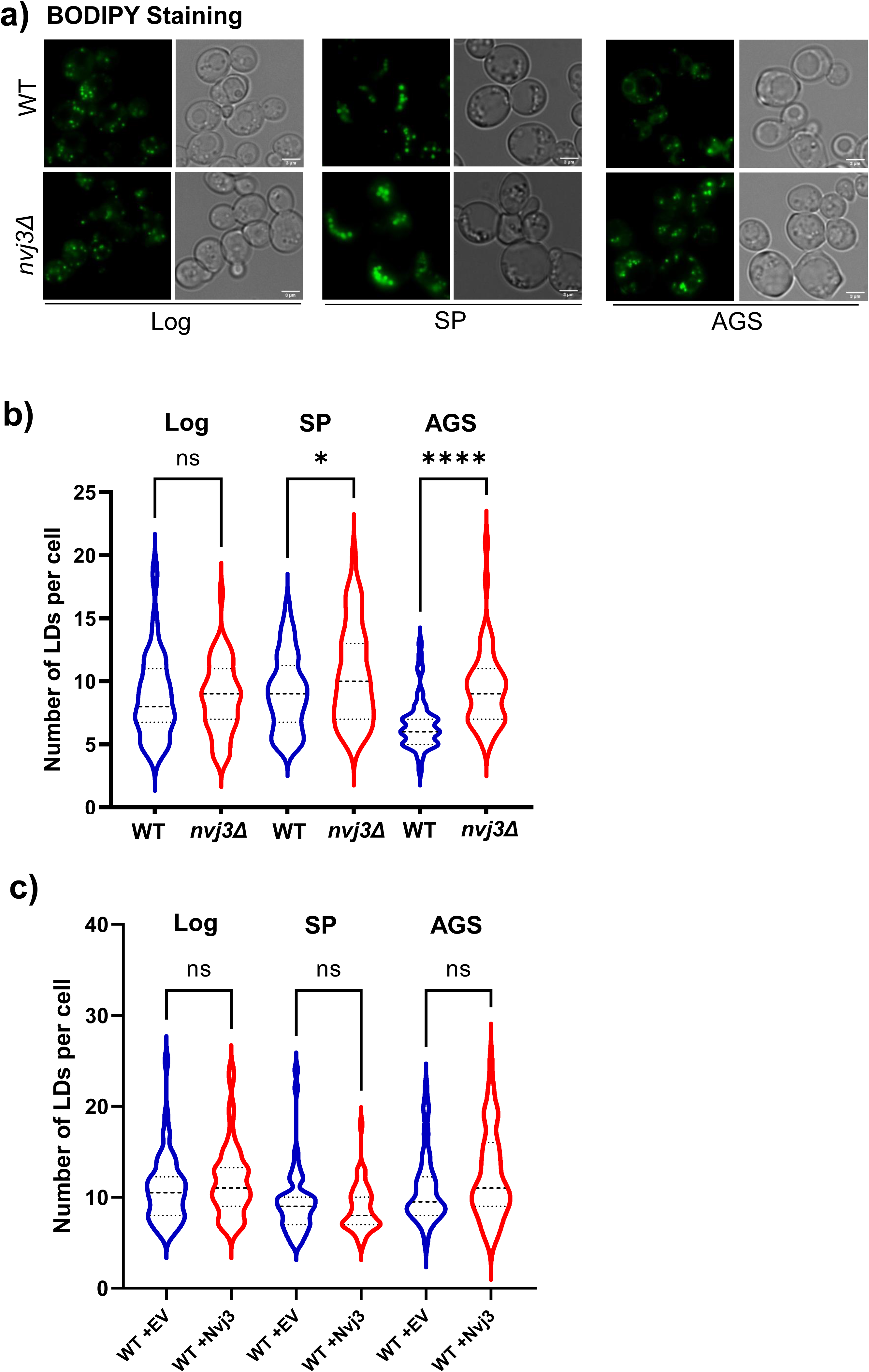
Nvj3 deletion causes LD accumulation during gradual and acute glucose starvation. A. Fluorescence images of BODIPY-stained LDs (green) in wild-type (WT) and *NVJ3* knock-out (*nvj3Δ*) yeast during log, stationary phase (48h), and acute glucose starvation (AGS; 0.001% glucose, 6h) and corresponding DIC images. Scale bar, 3 µm. B. Quantification of LD number per cell in WT and *nvj3Δ* strains at the indicated conditions (Log, SP, AGS). Data represent mean ± SD; *n* ≥ 100 cells per condition. *p* < 0.05 (*), **** *P* < 0.0001, ns = not significant (Student’s *t* test). C. Quantification of LDs per cell in WT cells expressing either an empty vector (EV) or *NVJ3* during Log, SP, and AGS. Mean ± SD; *n* ≥ 100 cells; ns = not significant.

We next assessed lipid composition by thin-layer chromatography (TLC) across growth and starvation conditions to determine how loss of Nvj3 impacts neutral lipid and phospholipid profiles (Figure 3). In log phase growth, *nvj3Δ* cells exhibited increases in TAG, sterol esters (SE), and FFAs relative to wild-type, indicating altered baseline lipid homeostasis (Figure 3a, Figure S2a). Upon entering SP, wild-type cells showed an increase in TAG and SE accompanied by reduction in FFAs, consistent with increased lipid storage during gradual nutrient depletion (Figure 3a). *nvj3Δ* cells displayed a broadly similar trend during SP, although neutral lipid levels (TAG and FFA) remained elevated relative to wild-type (Figure 3a).

**Figure 3.**
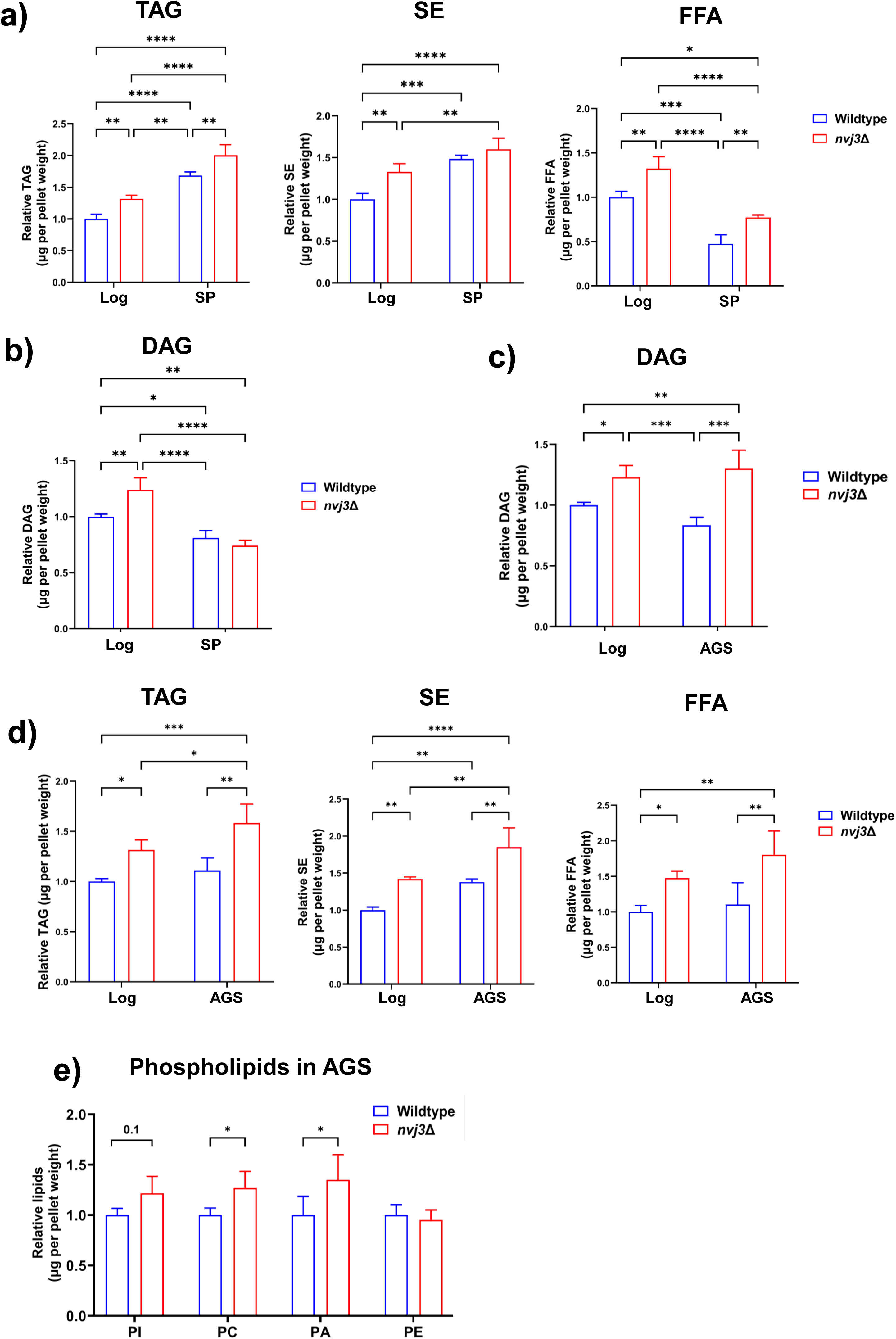
Loss of Nvj3 alters neutral and phospholipid composition under nutrient stress. A) Quantification of TAG (triacylglycerol), SE (sterol esters), and FFA (free fatty acids) levels from wild-type and *nvj3Δ* cells grown to log or SP (72h) measured by thin-layer chromatography (TLC). B) Quantification of DAG (diacylglycerol) levels in wild-type and *nvj3Δ* cells grown to log or SP (72h) measured TLC. C) Quantification of DAG levels in wild-type and *nvj3Δ* cells grown to log or AGS (acute glucose starvation) (72h) measured by TLC. D) Quantification of TAG, SE, FFA from wild-type and *nvj3Δ* cells grown to log or AGS measured by TLC. E) Quantification of phospholipid species phosphatidic acid (PA), phosphatidylethanolamine (PE), phosphatidylcholine (PC), and phosphatidylinositol (PI) from wild-type and *nvj3Δ* cells grown in AGS measured by TLC. Mean ± SD from three biological replicates. *P* < 0.05 (\**), < 0.01 (*\*\**), < 0.001 (*\*\*\*), < 0.0001 (****); Two-way ANOVA.

We next examined DAG, a central intermediate linking neutral lipid storage and phospholipid synthesis. In log phase cultures, DAG levels were elevated in *nvj3Δ* cells compared to wild-type (Figure 3b). Notably, this difference diminished upon entry into SP, as DAG levels in *nvj3Δ* cells declined toward wild-type levels (Figure 3b), suggesting that compensatory mechanisms activated during gradual starvation can partially normalize DAG homeostasis. In contrast, AGS elicited a distinct lipid signature where DAG levels remained elevated in *nvj3Δ*. Additionally, under AGS, *nvj3Δ* cells accumulated excess TAG, SE, and FFAs relative to wild-type (Figure 3c,d, S2a). Ergosterol (Erg) levels displayed a similar pattern to DAG in both starvation conditions, where elevated Erg levels in *nvj3Δ* are normalized during SP growth and not AGS (Figure S2a, b). Thus, accumulation of DAG (and Erg) was selective for acute, but not gradual, nutrient stress.

Because DAG serves as a critical branching point between TAG synthesis and phospholipid production, we then assessed phospholipid composition to determine how DAG accumulation alters phospholipid profile. Under AGS, *nvj3Δ* cells exhibited increased levels of phospholipids specifically PA and PC relative to wild-type (Figure 3e; Figure S2c). In contrast, phospholipid levels in log phase cultures were largely similar between genotypes, with the exception of increased PI in *nvj3Δ* cells (Figure S2c, d).

Together, these data indicate that Nvj3 contributes to baseline lipid homeostasis during growth. DAG accumulation under AGS suggests that loss of Nvj3 compromises efficient DAG utilization and lipid remodeling under conditions requiring rapid metabolic adaptation.

Because increased LDs and neutral lipids can arise either from enhanced synthesis or impaired degradation, we next examined whether loss of Nvj3 impairs LD breakdown.

### LD degradation pathways remain intact in *nvj3*Δ cells

To determine whether the increased LDs in *nvj3*Δ results from impaired breakdown, we assayed cytosolic lipolysis and vacuole-mediated lipophagy. For cytosolic TAG lipolysis, we used a cerulenin-based assay to block *de novo* FA synthesis, forcing cells to utilize TAG-derived FAs (Speer et al., 2023). Yeast contain three major TAG lipases, Tgl3, Tgl4, and Tgl5, of which Tgl3 performs the majority of TAG lipolysis (Kurat et al., 2006; Athenstaedt and Daum, 2003). TAG levels were measured in wild-type, *nvj3*Δ, and *tgl3*Δ yeast before (T_0_) and after 6 hours of treatment (T_6_). Cells were collected at early SP, when baseline TAG levels were comparable across wild-type and *nvj3*Δ, and elevated as expected in *tgl3*Δ strains (Figure 4a, b). TAG depletion kinetics were indistinguishable between wild-type and *nvj3*Δ cells, with both retaining ∼60% of TAG after 6 hours of cerulenin treatment, whereas *tgl3*Δ cells exhibited the expected defect retaining ∼80% of their TAG (Figure 4a, c). This data indicates that deletion of Nvj3 does not impair cytosolic TAG lipolysis.

**Figure 4.**
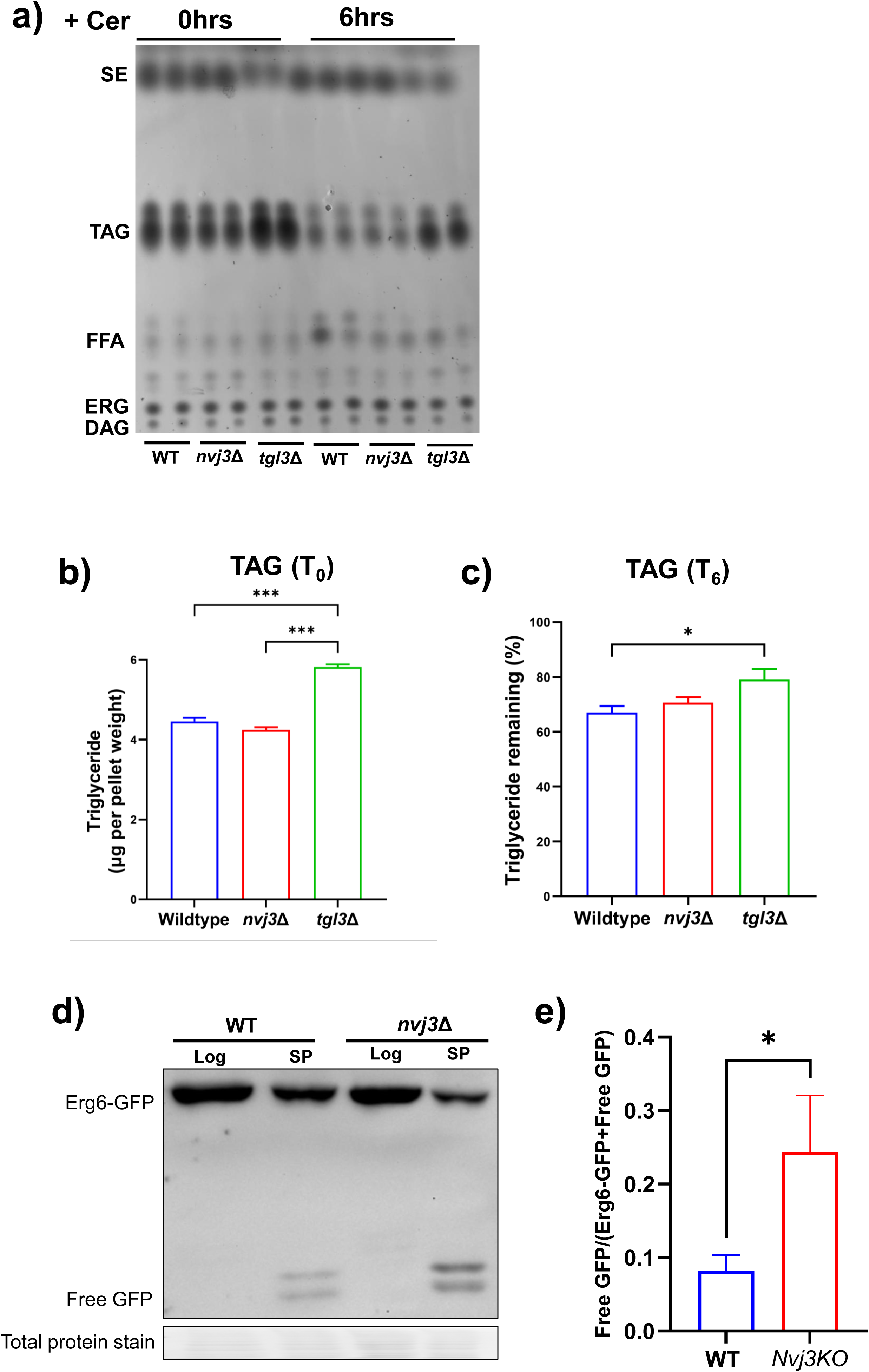
Lipolysis and lipophagy are not impaired in *nvj3Δ* cells. A) Thin-layer chromatography (TLC) of neutral lipid species from wild-type (WT), *nvj3Δ*, and *tgl3Δ* cells treated with cerulenin (10 µg/ml) for 0h and 6h. Bands corresponding to sterol esters (SE), triacylglycerols (TAG), free fatty acids (FFA), ergosterol (ERG), and diacylglycerols (DAG) are indicated. B) Quantification of TAG levels at 0h showing comparable lipid stores among WT and *nvj3Δ* strain prior to treatment. Mean ± SD, *n*=2. *P < 0.001 (*\*\*\*); One-way ANOVA. C) Quantification of percentage of TAG remaining after 6h of cerulenin treatment. WT and *nvj3Δ* cells show ∼60% TAG remaining, whereas *tgl3Δ* retains more TAG (∼80% remaining). *P* < 0.05 (*); One-way ANOVA. D) Western blot analysis of Erg6-GFP cleavage in WT and *nvj3Δ* during lipophagy induction at SP (72h). Free GFP bands released upon vacuolar degradation are indicated. Total protein staining is used as a loading control. E) Quantification of Erg6-GFP degradation from (D), comparing the extent of GFP cleavage between WT and *nvj3Δ.* Mean ± SD from three biological replicates.

Next, we asked whether increased LD abundance in *nvj3*Δ cells reflects impaired LD turnover by lipophagy. To assess lipophagy, we monitored vacuole-dependent degradation of LDs using Erg6-GFP cleavage assay, in which delivery of LD-associated Erg6-GFP to the vacuole results in proteolytic release of free GFP that can be detected by immunoblotting (van Zutphen et al., 2014). After prolonged SP growth (72h), *nvj3*Δ cells showed increased levels of free GFP compared to wild-type, indicating enhanced lipophagic flux under chronic starvation (Figure 4d, e).

After 6 hours of AGS, a time point at which LD accumulation was prominent in *nvj3*Δ cells, fluorescent imaging of BODIPY-stained LDs and FM4-64-labled vacuoles revealed no significant difference in LD-vacuole colocalization between wild-type and *nvj3*Δ cells (Figure S3a, b). This is consistent with prior work showing that lipophagy is minimal during early AGS and becomes more robustly activated during prolonged nutrient limitation (Seo et al., 2017; Fairman and Ouimet, 2022).

To further test whether LD turnover pathways remain functional in the absence of Nvj3, we induced lipophagy using nitrogen starvation, a condition also known to strongly activate vacuolar degradation of LDs (Klionsky et al., 2016). Under these conditions, *nvj3*Δ cells again displayed elevated Erg6-GFP cleavage relative to wild-type, confirming that lipophagic machinery is intact, and if anything, hyperactivated in the absence of Nvj3 (Figure S3c, d).

Together, the data indicate that increased LD abundance in *nvj3*Δ cells does not result from impaired LD degradation. Instead, LD turnover pathways remain functional across multiple starvation conditions, supporting the conclusion that LD accumulation in *nvj3Δ* cells reflects altered neutral lipid synthesis or metabolic routing upstream of degradation. Accordingly, we next tested whether Nvj3 regulates *de novo* TAG synthesis directly using a controlled LD biogenesis system.

### Nvj3 promotes efficient TAG synthesis under acute lipid induction

To examine how Nvj3 affects TAG synthesis, we generated yeast strains in which neutral lipid production could be tightly controlled (Figure 5a, d). This system eliminates sterol ester synthesis enzymes (*ARE1ΔARE2Δ*) and restricts TAG synthesis to a single inducible enzyme (Dga1 or Lro1) that is expressed from a galactose-inducible promoter. We refer to these strains as ΔLD. Therefore, in glucose, TAG synthesis is suppressed, while galactose addition acutely induces TAG production (Oelkers et al., 2002; Cartwright et al., 2015).

**Figure 5.**
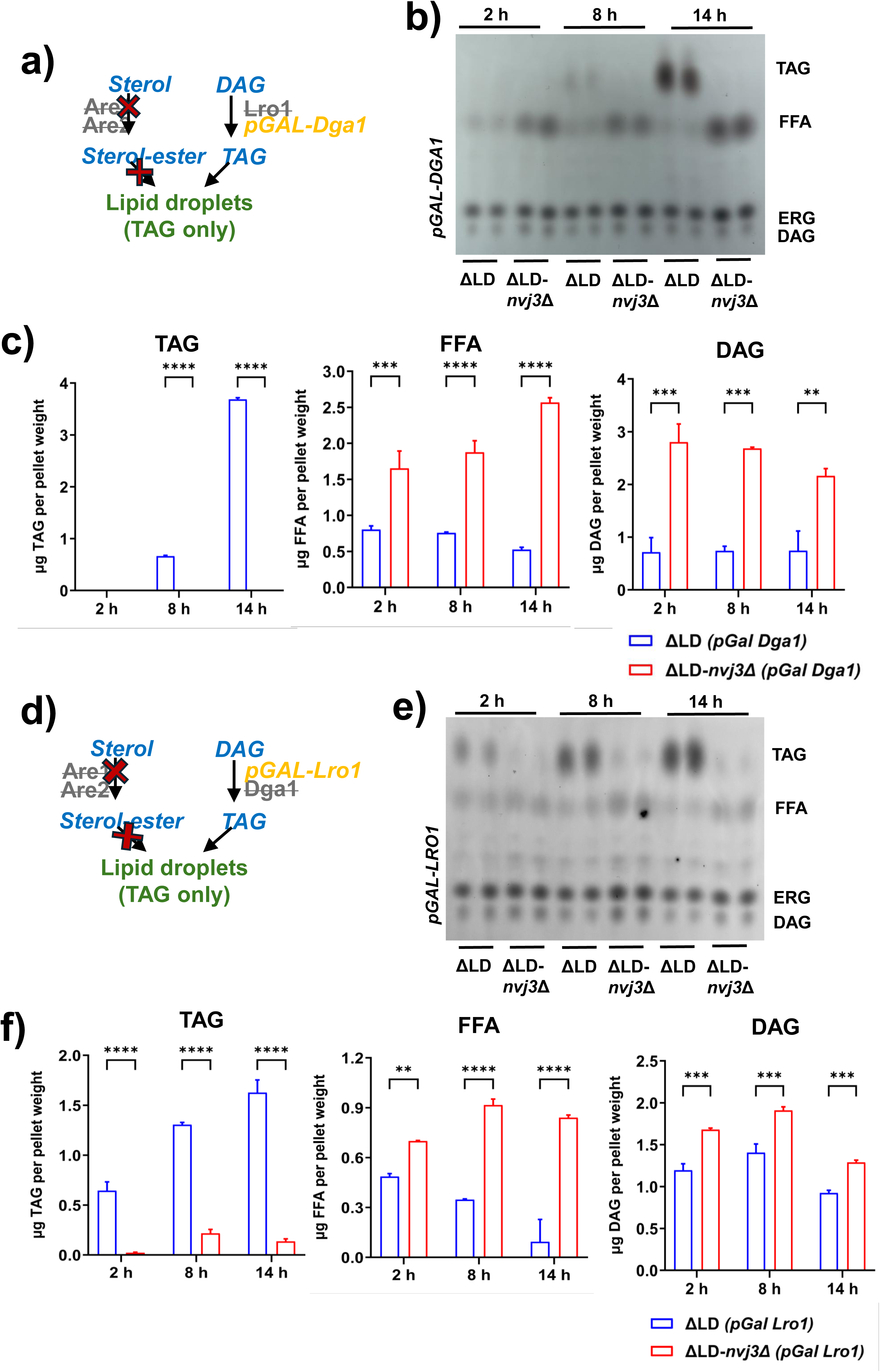
Nvj3 is required for efficient TAG synthesis during acute induction of Dga1– and Lro1-dependent pathways. A) Schematic of the inducible *pGAL-DGA1* system in *are1Δ are2Δ lro1Δ* yeast (ΔLD). In glucose, TAG synthesis is suppressed; upon galactose addition, Dga1 catalyzes acyl-CoA-dependent triacyglycerol (TAG) synthesis from diacylglycerol (DAG), enabling LD formation in the absence of sterol ester (SE) synthesis. B) Thin-layer chromatography (TLC) of neutral lipid species from *pGAL-DGA1* (ΔLD) and *pGAL-DGA1-nvj3Δ* (ΔLD-nvj3Δ) cells collected after 2, 8, and 14h of galactose induction. Bands corresponding to TAG, free fatty acids (FFA), ergosterol (ERG), and DAG are indicated. C) Quantification of TAG, DAG, and FFA in (b). *nvj3Δ* cells show strong accumulation of FFA and DAG with an almost complete block in TAG synthesis throughout the induction time course. D) Schematic of the inducible *pGAL-LRO1* system in *are1Δ are2Δ dga1Δ* yeast (ΔLD). Upon galactose induction, Lro1 catalyzes the conversion of diacylglycerol DAG and phospholipid acyl chains into TAG, allowing LD formation in the absence of SE synthesis. E) Thin-layer chromatography (TLC) of neutral lipid species from *pGAL-LRO1* (ΔLD) and *pGAL-LRO1-nvj3Δ* (ΔLD-*nvj3*Δ) cells harvested after 2, 8, and 14h of galactose induction. Bands corresponding to TAG, FFA, ERG, and DAG are indicated. F) Quantification of TAG and FFA levels from (e). ΔLD-*nvj3*Δ cells fail to accumulate TAG and instead accumulate FFA throughout the induction time course, indicating impaired Lro1-dependent TAG synthesis. Mean ± SD, n = 2. *P < 0.01 (**), < 0.001 (***), < 0.0001 (****)*; Two-way ANOVA.

We next performed a kinetic assay by switching pGAL-DGA1 ΔLD yeast from glucose to galactose-containing media and tracking TAG synthesis over time using TLC (2, 8, and 14 hours). Following galactose induction, pGAL-DGA1 ΔLD cells displayed a progressive decline in FFAs accompanied by robust TAG accumulation, reflecting efficient conversion of FFAs into TAG (Figure 5b, c). In contrast, pGAL-DGA1 ΔLD-*nvj3Δ* cells exhibited persistent FFA and DAG accumulation and a pronounced delay in TAG synthesis across the induction time course (2-14 hours), indicating a defect in TAG synthesis when Nvj3 is absent (Figure 5b, c).

To determine whether this defect was specific to the acyl-CoA-dependent TAG synthase Dga1, we performed parallel experiments using a galactose-inducible *LRO1* strain lacking Dga1 and sterol ester synthases (pGAL-LRO1; Figure 5d). Similar to the Dga1-inducible system, pGAL-LRO1 ΔLD-*nvj3Δ* cells showed delayed TAG accumulation and elevated FFAs and DAG relative to pGAL-LRO1 ΔLD cells (Figure 5e, f), demonstrating that Nvj3 is required for efficient TAG synthesis at early induction time points through both Dga1-and Lro1-mediated pathways.

To determine how the delay in inducible TAG synthesis affects phospholipid levels, we measured phospholipid composition during DGA1 and LRO1 induction using TLC. In pGAL-DGA1 ΔLD-*nvj3*Δ cells, PA levels were significantly less as early as 2 hours after induction compared to pGAL-DGA1 ΔLD cells (Figure S4a, b). Simultaneously, PI levels increased progressively only in pGAL-DGA1 ΔLD-*nvj3*Δ cells and remained stable in pGAL-DGA1 ΔLD cells (Figure S4c). PC and PE increased similarly in both strains with no significant differences (Figure S4c). A similar phospholipid pattern was observed in pGAL-LRO1 ΔLD*-nvj3*Δ; PA depletion accompanied by elevated PI (Figure S4d-f). PC was elevated in pGAL-LRO1 ΔLD*-nvj3*Δ at 8h and PE was trending down with no significant change between strains (Figure S4d-f). Together, this data suggest that in the absence of Nvj3, PA and DAG are preferentially routed towards phospholipid pathways rather than efficiently channeled toward TAG synthesis.

### Nvj3 couples TAG synthesis to efficient LD formation

We next examined how impaired TAG synthesis and phospholipid remodeling in ΔLD*-nvj3*Δ cells affect LD biogenesis by monitoring LD formation using BODIPY staining in the controlled TAG induction system. Upon galactose induction of pGAL-DGA1, ΔLD control cells formed visible LDs within 2-4 hours, with LD number increasing progressively through 16 hours, consistent with efficient TAG synthesis and LD assembly (Figure 6a; Figure S5a). In contrast, pGAL-DGA1 ΔLD-*nvj3Δ* cells exhibited a pronounced delay in LD appearance, with minimal or no detectable LDs at early time points (Figure 6a; Figure S5a). Quantification confirmed a persistent reduction in LD number per cell across the induction time course (Figure 6c; Figure S5b).

**Figure 6.**
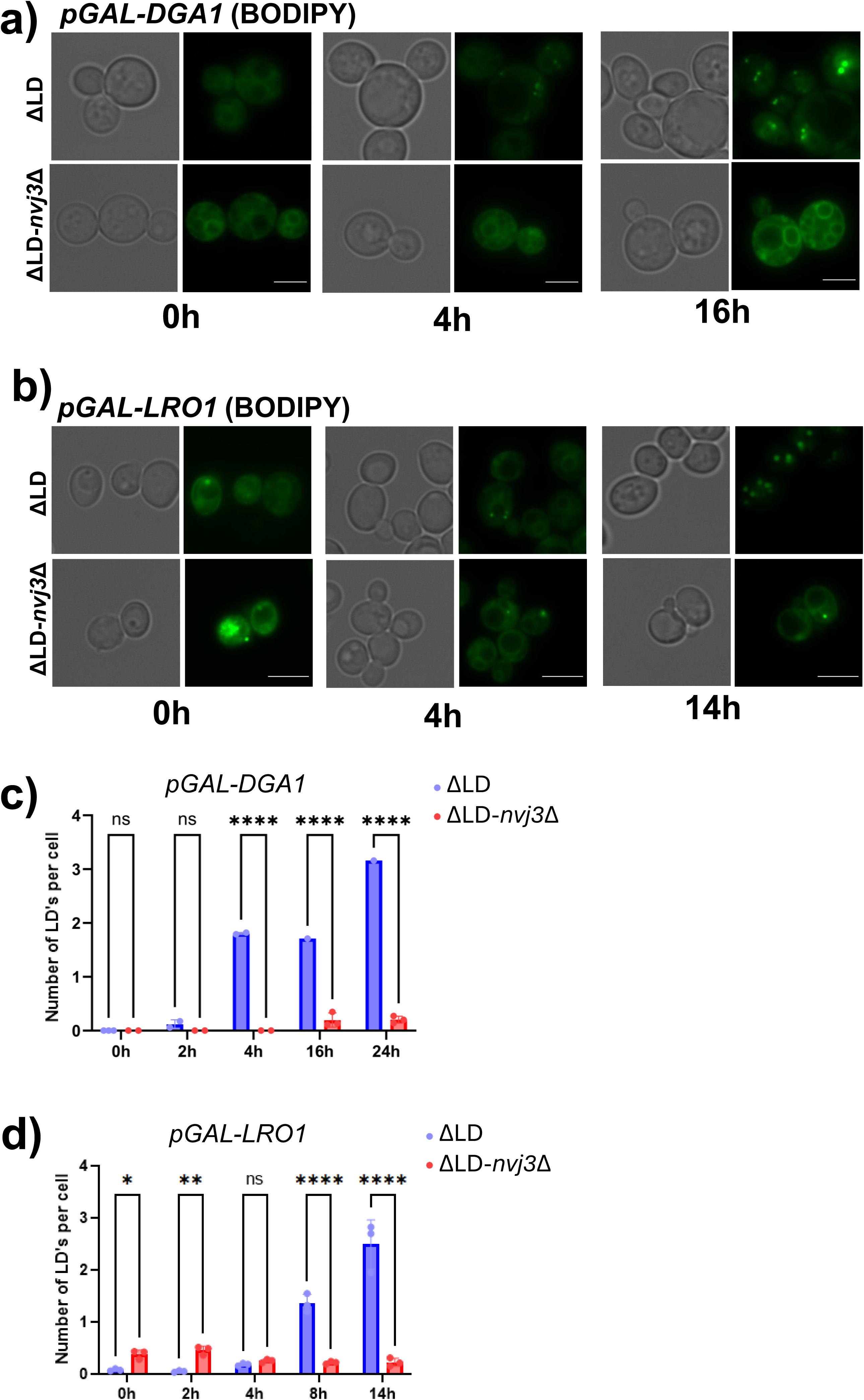
Nvj3 is required for Lro1-dependent TAG synthesis and lipid droplet formation. A) Fluorescence micrographs of BODIPY staining at 0, 4, and 16h post-galactose induction in *pGAL-DGA1* (ΔLD) and *pGAL-DGA1-nvj3Δ* (ΔLD-*nvj3*Δ). B) Fluorescence micrographs of BODIPY staining at 0, 4, 14h post-galactose induction in *pGAL-LRO1* (ΔLD) and *pGAL-LRO1-nvj3Δ* (ΔLD-*nvj3*Δ). Scale bar, 3 µm. C) Quantification of average number of LD per cell from (a). *pGAL-DGA1-nvj3Δ* (ΔLD-nvj3Δ) cells contain significantly fewer LDs at all time points. D) Quantification of average number of LD per cell from (a). *pGAL-LRO1-nvj3Δ* (ΔLD-nvj3Δ) cells contain significantly fewer LDs at all time points. Mean ± SD, *n* ≥ 100 cells; *P < 0.05 (*), < 0.01 (**), < 0.0001 (****),* ns = not significant; Two-way ANOVA.

At prolonged induction time points (16-24h), BODIPY signal became detectable in pGAL-DGA1 ΔLD-*nvj3Δ* cells; however, this signal was largely diffuse and ER-associated rather than forming discrete punctate LDs, in striking contrast to pGAL-DGA1 ΔLD controls (Figure 6a, Figure S5c). Thus, although neutral lipid synthesis eventually occurs in the absence of Nvj3, TAG fails to be efficiently packaged into LDs.

Notably, the defect was more severe than that previously observed in pGAL-DGA1-*mdm1*Δ, which retain the ability to form detectable LDs by 6 hours of induction (Hariri et al., 2019). To enable direct comparison, we repeated the pGAL-DGA1 induction in *mdm1*Δ and *nvj3*Δ backgrounds and tracked LD formation at 4h and 6h using BODIPY staining. Whereas ΔLD*-mdm1*Δ cells retained the ability to generate a limited number of LDs, ΔLD*-nvj3*Δ cells showed near complete block in assembling discrete LDs with markedly reduced LD numbers and low fraction of cells containing LDs (Figure S5e, f). Thus, loss of Nvj3 is more detrimental to Dga1-driven LD assembly than loss of Mdm1, indicating that Nvj3 plays a critical role in enabling TAG packaging in LDs downstream of Dga1 activity.

We next examined LD formation in the pGAL-LRO1 induction system, where Lro1 is the sole TAG synthase. In contrast to the Dga1-driven system, pGAL-LRO1 ΔLD-*nvj3Δ* cells formed detectable punctate LDs as early as 2 hours post-inductions (Figure 6b; Figure S5c). Although the LD numbers in pGAL-LRO1 ΔLD-*nvj3Δ* did not increase substantially over time, they remained consistently lower than pGAL-LRO1 ΔLD controls throughout the induction time course (Figure 6b, d; Figure S5c, d).

Interestingly, LD formation in the Lro1-driven ΔLD-*nvj3Δ* background was quantitatively more efficient that in the Dga1-driven system (Figure 6c, d; S5b, d). At equivalent induction times, Lro1-induced ΔLD-*nvj3Δ* cells displayed clear punctate LDs and reduced diffuse, ER-associated BODIPY signal compared to Dga1-induced ΔLD-*nvj3Δ* cells (Figure 6a, b). These observations indicate that while Nvj3 contributes to optimal LD production in both pathways, Lro1-mediated TAG synthesis is less dependent on Nvj3 for initial coupling of TAG synthesis to LD assembly.

The emergence of diffuse, ER-associated neutral lipid signal in the Dga1-driven system prompted us to test whether prolonged TAG synthesis could overcome this defect, leading us to examine TAG accumulation and LD morphology under extended induction conditions.

### Loss of Nvj3 uncouples Dga1-driven TAG synthesis from LD formation

To determine whether extended TAG synthesis could overcome the defect in LD formation observed in Dga1-inducible ΔLD-*nvj3Δ* cells, we extended galactose induction to 48 hours and assessed neutral lipid accumulation and LD morphology by TLC and BODIPY staining, respectively. Under these conditions, pGAL-DGA1 ΔLD-*nvj3Δ* cells accumulated substantial amounts of TAG, reaching levels that exceeded those observed in the pGAL-LRO1 ΔLD-*nvj3Δ* system at comparable time points (Figure 7a, b). Thus, loss of Nvj3 does not completely impair the capacity for TAG synthesis *per se* during prolonged induction.

**Figure 7.**
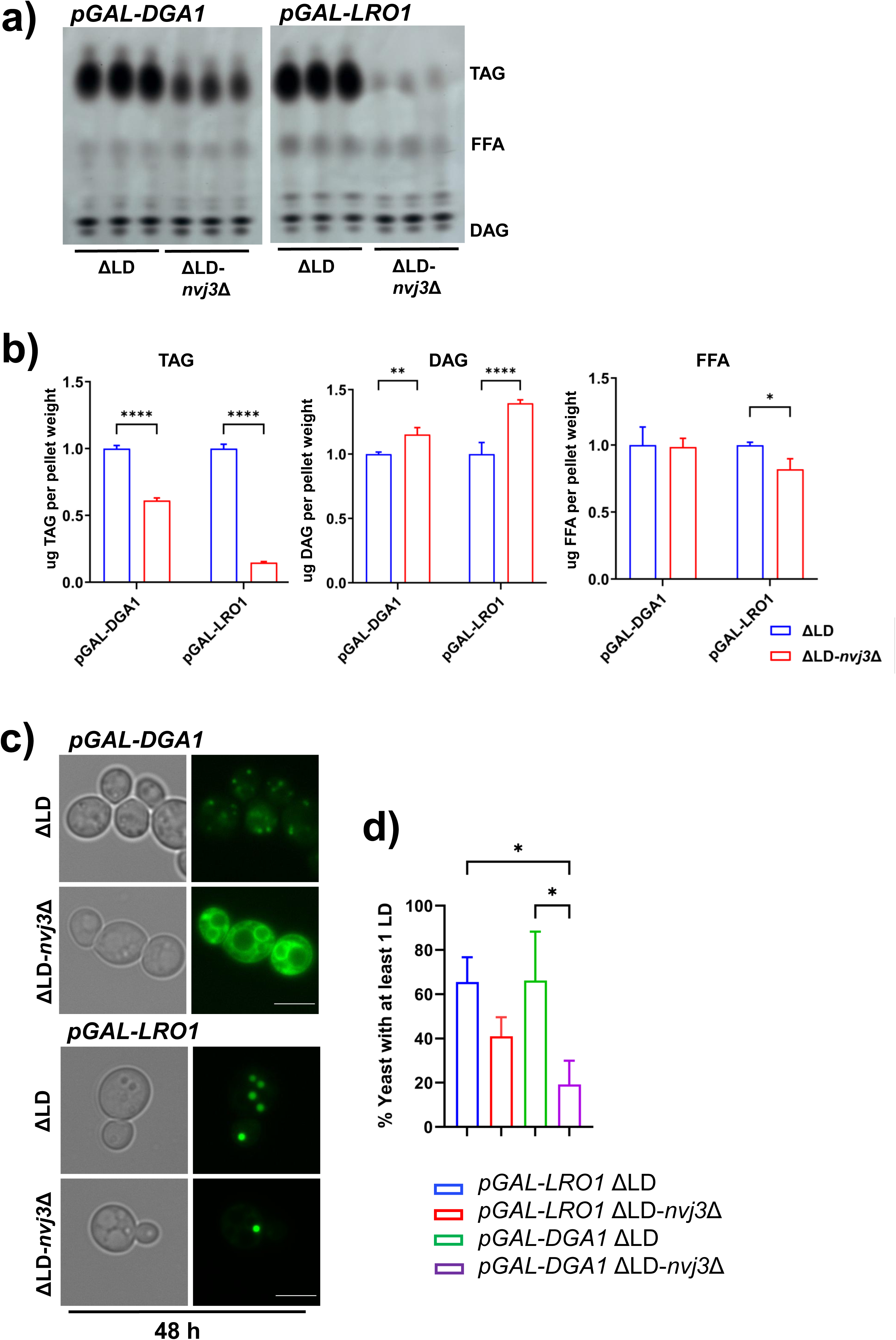
Nvj3 is required for both Dga1– and Lro1-dependent triacylglycerol synthesis and lipid droplet formation. A) Thin-layer chromatography (TLC) of neutral lipids from *pGAL-DGA1* and *pGAL-LRO1* cells lacking Are1, Are2, and Lro1 or Dga1 (ΔLD) and their *nvj3*Δ counterparts (ΔLD-*nvj3*Δ) after 48h galactose induction. Bands corresponding to triacylglycerol (TAG), free fatty acid (FFA), and diacylglycerol (DAG) are indicated. B) Quantification of neutral lipid classes from (a). *nvj3*Δ cells show significantly reduced TAG accumulation with concomitant DAG and FFA increases in both pGAL-DGA1 and pGAL-LRO1 backgrounds. Mean ± SD, n = 3; *P < 0.05 (*), < 0.01 (**), < 0.0001 (****);* Two-way ANOVA. C) Fluorescence micrographs of BODIPY stain in *pGAL-DGA1* and *pGAL-LRO1* cells stained with BODIPY after 48h galactose induction. *nvj3*Δ cells exhibit defective LD formation under both Dga1– and Lro1-driven TAG synthesis. Scale bar, 3 µm. D) Quantification of imaging data in (c); percentage of cells containing ≥1 LD. Loss of Nvj3 significantly reduces LD biogenesis in both Dga1– and Lro1-dependent systems. Mean ± SD, *n* ≥ 100 cells; *P < 0.05 (*), < 0.01 (**), < 0.0001 (****);* One-way ANOVA.

Despite this TAG accumulation, pGAL-DGA1-*nvj3Δ* cells failed to form discrete LDs at 48h. Instead, BODIPY staining revealed predominantly diffuse, ER-associated neutral lipid signal, in stark contrast to ΔLD controls, which displayed abundant punctate LDs at 48 hours (Figure 7c, d). This phenotype indicates a persistent defect in LD assembly rather than a simple kinetic delay.

In contrast, pGAL-LRO1 ΔLD-*nvj3Δ* cells formed distinct, punctate LDs by 48 hours, despite overall lower TAG accumulation relative to the Dga1-driven system (Figure 7a-d). In these cells, BODIPY signal was largely confined to LDs, with nondetectable ER-associated neutral lipid signal. Thus, while Lro1-mediated TAG synthesis is delayed in the absence of Nvj3, these cells remain capable of supporting efficient LD formation.

Together, these results demonstrate that TAG accumulation and LD formation can be uncoupled in the absence of Nvj3, particularly in the context of Dga1-driven TAG synthesis. Although Dga1-driven TAG can accumulate to relatively high levels, this TAG remains inefficiently packaged into LDs, instead persists in a diffuse ER-associated state in absence Nvj3. These findings suggest that Nvj3 is required not simply for TAG production per se, but rather for organizing the lipid environment in which TAG is efficiently converted into discrete LDs. We therefore next examined whether loss of Nvj3 alters the subcellular distribution of DAG, and the abundance of Dga1.

### Nvj3 maintains DAG localization and supports Dga1 expression

The inducible TAG synthesis experiments revealed that loss of Nvj3 uncouples TAG accumulation from efficient LD formation, particularly in the context of Dga1-driven TAG synthesis. However, under steady state *nvj3Δ* cells ultimately accumulate TAG and LDs, suggesting that loss of Nvj3 does not block TAG synthesis *per se* but may instead regulate upstream lipid organization that determines how TAG synthesis is translated into productive LD assembly over time. We therefore examined whether loss of Nvj3 alters the subcellular distribution of DAG and the expression and recruitment of Dga1 under non-inducible, steady state conditions.

To assess the spatial distribution of DAG, we visualized DAG using a previously characterized fluorescent C1-domain–based ER-DAG sensor derived from Protein Kinase D (Choudhary et al., 2018). In wild-type cells under AGS, DAG signal was predominantly associated with ER membranes. In contrast, *nvj3Δ* cells showed reduced ER-associated DAG signal accompanied by a marked increase in DAG sensor signal within the vacuole (Figure 8a, b). Supplementation with OA which promotes neutral lipid synthesis and LD expansion, increased punctate DAG signal in wild-type cells but failed to restore ER-associated DAG in *nvj3Δ* cells where the signal remained predominantly in the vacuole (Figure S6a, b). These data indicate that Nvj3 is required to maintain proper DAG partitioning during metabolic stress.

**Figure 8.**
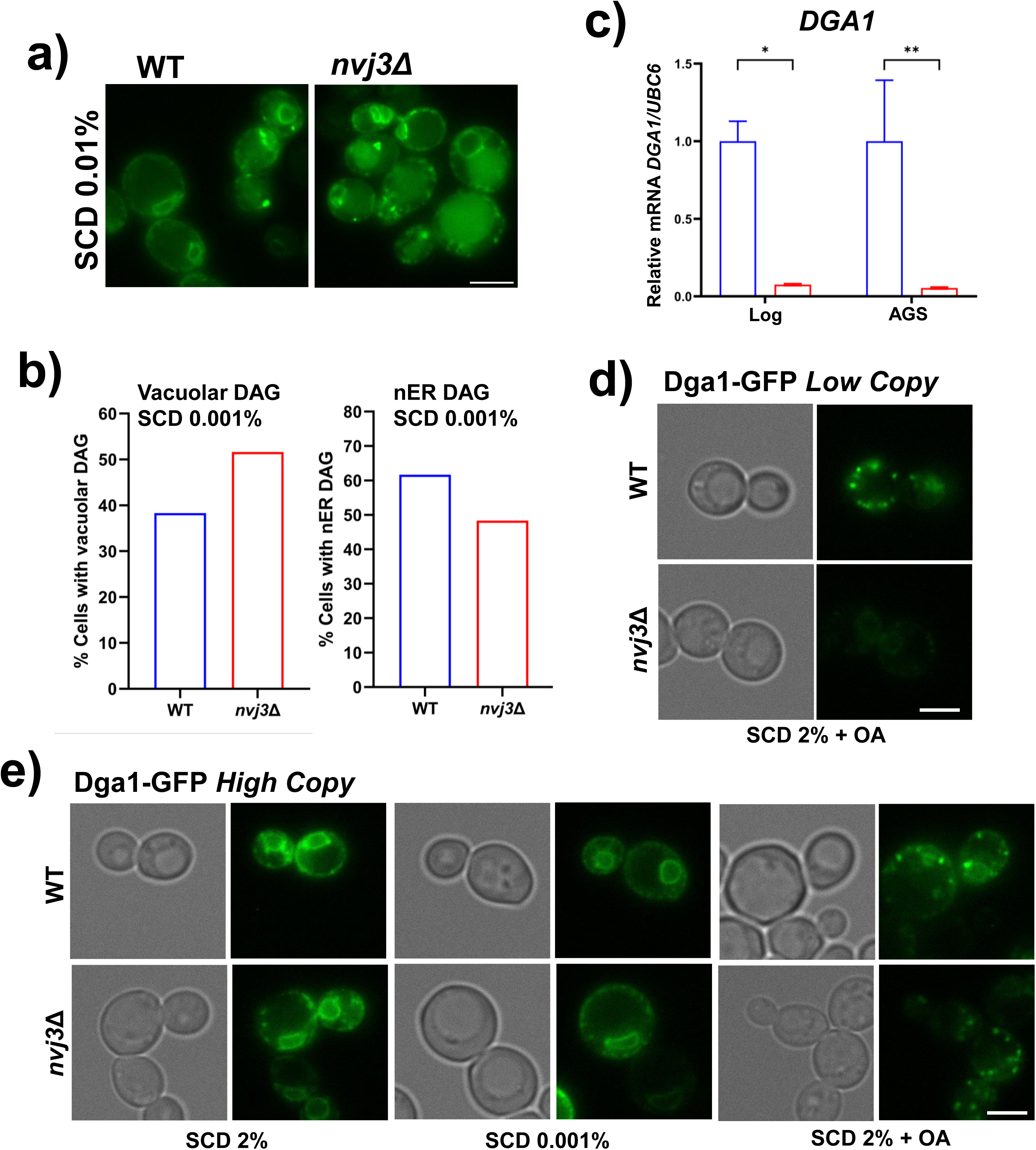
Loss of Nvj3 alters DAG localization and reduces Dga1 abundance. a) Fluorescent imaging of GFP-tagged DAG sensor (C1-domain from Protein Kinase D) in wild-type (WT) and *nvj3*Δ cells grown under low glucose (SCD 0.01%). b) Quantification of (a) DAG signal localization (percent cells). *nvj3*Δ cells display increased DAG retention at the vacuole (vacuolar DAG) and reduced nER-associated DAG, *n* ≥ 100 cells. c) Relative mRNA expression of *DGA1* in wild-type and *nvj3*Δ cells during log phase and acute starvation. Mean ± SEM, n = 4; *P < 0.05 (*), < 0.01 (**);* Two-way ANOVA. d) Fluorescent imaging of Dga1-GFP overexpression via low-copy plasmid in wild-type and *nvj3*Δ cells grown in oleate-supplemented (SCD 2% + OA) conditions. e) Fluorescent imaging of Dga1-GFP overexpression via high-copy plasmid in wild-type and *nvj3*Δ cells grown in rich (SCD 2%), low-carbon (SCD 0.001%), or oleate-supplemented (SCD 2% + OA) conditions. Scale bar, 3 µm.

Because DAG plays as a central role as a precursor of TAG synthesis, we next tested whether altered organization DAG in *nvj3Δ* cells could also impact TAG the abundance and recruitment of the TAG synthase Dga1. We first attempted to tag endogenously tag Dga1 with a fluorescent marker but were unable to obtain detectable signal only in the Nvj3 deletion strains, consistent with very low Dag1 abundance under this condition. Consistent with this, quantitative RT-PCR revealed significantly reduced *DGA1* transcript levels in *nvj3Δ* cells both in log phase and AGS conditions, whereas *LRO1* transcript levels were unchanged (Figure 8c, S6c).

We therefore used a plasmid-based expression to visualize Dga1 localization by live-cell fluorescent imaging under nutrient-replete conditions (SCD2%), AGS (SCD, 0.001%), and OA supplementation (SCD2%+OA), which robustly stimulates neutral lipid synthesis, LD expansion, and Dga1 recruitment to LD-associated domains. With a low-copy plasmid, Dga1-GFP was readily detectable in wild-type cells and showed punctate localization pattern that was particularly dominant upon OA supplementation (Figure 8d). In contrast, Dga1-GFP signal in *nvj3Δ* cells was markedly reduced under nutrient-replete and AGS and remained comparatively weak following OA supplementation, consistent with the conclusion that Dga1 abundance is diminished when Nvj3 is absent (Figure 8d, S6d).

To distinguish reduced abundance from a defect in targeting, we expressed Dga1-GFP from a high-copy plasmid. High-copy expression restored Dga1-GFP signal and punctate localization in *nvj3Δ* cells across all conditions (Figure 8e), demonstrating that Dga1 retains its intrinsic ability to associate with ER/LD domains when sufficiently expressed. Together, these data support a model in which Nvj3 promotes proper DAG partitioning during stress and also supports Dga1 abundance, while Dga1 recruitment/localization per se is not fundamentally impaired in the absence of Nvj3.

Finally, to assess the contributions of Dga1 and Lro1 to LD accumulation in *nvj3Δ* at cells in steady state, we performed genetic epistasis analysis using Erg6-GFP as an LD marker under AGS conditions. Consistent with our imaging and lipid profiling data, *nvj3Δ* cells exhibited elevated Erg6-GFP signal compared to wild-type, reflecting increased LD content (Figure S6e, f). Deletion of *DGA1* in the *nvj3Δ* background significantly reduced Erg6-GFP signal toward wild-type levels, whereas deletion of *LRO1* displayed no similar reduction (Figure S6e, f). These results indicate that despite reduced Dga1 expression, Dag1-dependent TAG synthesis remains a major driver of LD accumulation in *nvj3Δ* cells during AGS, consistent with a model in which Nvj3 regulates both DAG distribution and Dga1 availability to promote efficient LD assembly.

## Discussion

Lipid droplet biogenesis is increasingly recognized as a spatially organized process that depends not only on TAG synthesis but also on the cellular and local context in which neutral lipids are produced (Choudhary et al., 2020; Olzmann and Carvalho, 2019; Hariri et al., 2018; Zoni et al., 2021; Henne et al., 2025). In this study, we identify Nvj3 as a stress-responsive regulator of TAG synthesis and LD formation during acute metabolic transitions. Our data support a model in which Nvj3 regulates the efficiency and timing of LD formation by influencing the availability and utilization of lipid intermediates during stress.

A central finding of this work is that TAG accumulation and LD formation can be experimentally uncoupled in *nvj3Δ* cells, particularly in the context of Dga1-dependent TAG synthesis (Figure 6a, 7c). Using inducible TAG synthesis systems, we show that loss of Nvj3 causes a pronounced delay in TAG accumulation and LD formation during acute induction (Figure 5a-c; Figure 6a, c). Importantly, this phenotype does not reflect a complete block in TAG synthesis as prolonged induction ultimately results in substantial TAG accumulation in *nvj3Δ* cells (Figure 7a, b). However, newly synthesized TAG is inefficiently packaged into discrete LDs and instead remains associated with ER membranes (Figure 7c). These data demonstrate that, under these conditions, bulk TAG production alone is insufficient to drive timely LD formation in Nvj3-deficient cells.

The pathway specificity of this phenotype provides important mechanistic insight. Although Lro1-mediated TAG synthesis is also delayed in *nvj3Δ* cells, LD formation proceeds more efficiently in this context compared to Dga1-driven synthesis (Figure 5d-f; Figure 6b, d). This distinction suggests that Dga1-dependent TAG synthesis is particularly sensitive to upstream constraints imposed by deletion of Nvj3, whereas luminal Lro1-mediated TAG synthesis can partially bypass these constraints. Thus, Nvj3 does not broadly “couple” TAG synthesis to LD formation, but instead, it modulates how efficiently specific TAG synthesis pathways are translated into LD assembly during acute stress.

What constraints does Nvj3 deletion impose upstream of TAG synthesis pathways? Loss of Nvj3 alters lipid profiles in a context-dependent manner. During acute induction of TAG synthesis, *nvj3Δ* cells show reduced TAG and PA alongside accumulation of DAG, PI, and FFAs (Figure 5, Figure S4). This pattern is consistent with inefficient routing of lipid intermediates toward TAG synthesis, with DAG and PA being diverted into phospholipid pathways, particularly PI. Accumulation of FFAs further suggests delayed or inefficient coupling FA activation and acylation, a known rate-limiting step for TAG synthesis that is regulated at the NVJ during stress (Hariri et al., 2019).

In contrast, under steady-state AGS, *nvj3Δ* cells accumulate TAG together with increased DAG, PA, PI, and FFAs, indicating that lipid synthesis pathways remain active and can eventually support TAG accumulation and LD formation (Figure c-e; Figure S2c, d). The divergence between inducible and steady state conditions suggest that prolonged growth without Nvj3 engages compensatory phospholipid remodeling pathways that are not accessible during acute metabolic transitions. Together, these data indicate that Nvj3 is not required to maintain global lipid synthesis capacity but instead promotes timely and efficient routing of lipid intermediates during rapid metabolic shifts, when compensatory pathways are insufficient.

What impacts the routing of DAG in *nvj3Δ* cells? Direct imaging using a fluorescent DAG sensor supports altered partitioning of DAG in the absence of Nvj3. In wild-type cells during AGS, DAG signal is enriched at ER membranes, consistent with its role as a substrate for TAG synthesis. In contrast, *nvj3Δ* cells show reduced ER-associated DAG signal and accumulation of a prominent DAG pool within the vacuole (Figure 8a, b). Although the origin of this vacuolar DAG remains unclear, it could potentially be explained by the enhanced lipophagy which is more prominent in *nvj3Δ* cells during prolonged starvation. Regardless of its source, DAG redistribution away from the ER in *nvj3Δ* cells is consistent with reduced accessibility of DAG at ER regions competent for LD formation, particularly during acute metabolic stress.

Although we do not directly demonstrate DAG transfer by Nvj3, its behavior and domain architecture suggest potential parallels with emerging lipid-transfer paradigms. Recent studies of Atg2 and related proteins have revealed extended hydrophobic conduits that enable bulk lipid movement between membranes during autophagosome expansion and LD growth (Elhan et al., 2025; Korfhage et al., 2025). These findings provide a conceptual framework for how Nvj3 might act, either independently or in cooperation with Mdm1 and other contact site components, to promote efficient lipid channeling or retention during stress. Whether Nvj3 directly transfers lipids, stabilizes lipid-enriched ER microdomains, or restricts diffusion of intermediates such as DAG away from LD-competent regions remains an important question for future work.

Loss of Nvj3 is also associated with reduced expression of *DGA1*, while *LRO1* transcript levels remain unchanged (Figure 8a; S6c). Imaging endogenously tagged Dga1-GFP was not feasible in the *nvj3Δ* background, consistent with low basal expression. However, high-copy expression of Dga1-GFP restores punctate localization and LD association in *nvj3Δ* cells, demonstrating that Nvj3 is not required for Dga1 targeting or membrane association per se. Notably, under oleate supplementation, which robustly promotes LD expansion and Dga1 recruitment, Dga1-GFP signal remains reduced in nvj3Δ cells realtive to wild type, suggesting that Nvj3 supports efficient accumulation or stabilization of Dga1 under lipid-loading conditions.

Although transcriptional regulation of DGA1 remains incompletely understood, its expression is known to be stress-responsive and sensitive to metabolic state (Oelkers et al., 2002; Sorger and Daum, 2002). Reduced Dga1 abundance in nvj3Δ cells likely exacerbates delayed TAG synthesis during acute stress but does not eliminate Dga1 function. Consistent with this, genetic epistasis analysis shows that Dga1 remains a major contributor to LD accumulation during AGS even in the absence of Nvj3, indicating that Nvj3 constrains the efficiency and timing of Dga1-dependent TAG utilization rather than its fundamental capacity.

Finally, the behavior of Nvj3 further highlights functional diversity within the SNX-RGS protein family (Hariri and Henne, 2022). Nvj3 requires Mdm1 for stable recruitment to the NVJ yet also associates with LDs independently of this tether (Henne et al., 2015). This dual localization suggests that distinct SNX-RGS proteins or heteromeric assemblies may act in different cellular contexts to reorganize lipid flux at membrane contact sites, facilitating lipid storage, membrane remodeling, or stress-induced membrane damage repair as needed. In this view, Nvj3 functions not as a dedicated LD biogenesis factor, but as a stress-adaptive organizer that fine tunes the efficiency with which lipid intermediates are routed into lipid synthesis pathways.

In summary, we propose that Nvj3 regulates the efficiency of TAG synthesis and LD formation during metabolic stress by shaping lipid intermediate availability and utilization. Loss of Nvj3 does not abolish TAG production or LD accumulation, but delays their productive integration during acute stress, particularly for Dga1-dependent pathways, revealing a previously unappreciated layer of spatial regulation in LD biogenesis. This work highlights how membrane contact site proteins can act as context-dependent modulators of lipid flux, enabling cells to adapt lipid pathways to rapidly changing metabolic demands. Future studies will address whether Nvj3 directly participates in lipid transfer, how it functionally and physically cooperates with Mdm1, and whether distinct SNX-RGS assemblies act at different contact sites to adapt lipid flux for storage, membrane remodeling, or stress-induced repair.

## Materials and methods

### Molecular biology, yeast genetics, and growth conditions

All yeast strains and plasmids used in this study are listed in Tables S1 and S2, respectively. Yeast genetic manipulations were performed using the standard lithium acetate transformation method. Unless otherwise noted, cells were grown in synthetic complete (SC) medium containing 2% dextrose at 30°C with shaking at 225 rpm. Log-phase cultures were obtained by growing cells to an optical density (OD_600_) of 0.8–1.0. All washes were performed with sterile, deionized water.

#### Starvation protocols

For acute starvation, log-phase cells grown in SC + 2% dextrose were collected, washed twice, and resuspended in SC containing 0.001% dextrose for 6 hr. For gradual starvation, cultures were grown in SC + 2% dextrose for the indicated number of hours without media replenishment. Where indicated, cerulenin (Sigma-Aldrich; 10 µg/mL) was added at 48h of gradual starvation.

#### Cell harvesting

For lipid analysis, cells were harvested by centrifugation at 4000 × g for 5 min, washed once with sterile water, and transferred to pre-weighed microcentrifuge tubes. Pellets were recentrifuged, the supernatant was removed, and wet weights were recorded. Cell pellets were flash-frozen in liquid nitrogen and stored at –80°C. A similar harvesting procedure was used for other downstream assays unless otherwise specified.

### Growth assays

Precultures were grown using SC media with 2% dextrose at standard growth conditions overnight. Precultures then were diluted to 0.05 OD_600_ in triplicate in a 96-well plate to a total volume of 200 µL, accounting for specified experimental conditions. Cell density was measured every 20 minutes using a SpectraMax i3x microplate reader at 30°C shaking at 225 rpm for 48 total hours. The resulting data was normalized against wells containing only liquid media to account for background absorbance and graphs generated using GraphPad Prism.

### qPCR

Cells were harvested by centrifugation (4000 × g, 5 min), the supernatant was aspirated, and pellets were flash-frozen in liquid nitrogen before storage at −80 °C. After thawing on ice, total RNA was isolated using the NucleoSpin RNA Plus kit (Takara Bio USA, #740984) according to the manufacturer’s protocol. Briefly, pellets were resuspended in 350 µL LBP buffer and ∼0.5 volume of 0.5-mm acid-washed glass beads before mechanical lysis using a bead beater (4 × 1-min cycles at maximum speed with 1-min cooling intervals on ice). RNA was further purified using the same kit and quantified using a Nanodrop spectrophotometer. cDNA was synthesized using the PrimeScript RT reagent kit with genomic DNA eraser (Takara Bio USA, #RR092A). qPCR was performed in 10-µL reactions using PowerUp SYBR Green Master Mix (Thermo Fisher, #A25777). Transcript levels were normalized to the housekeeping genes *UBC6* or *TAF10*. Primer efficiencies (95–105%) were validated empirically, and all primer sequences are provided in Table S3.

### Light microscopy

Live-cell imaging of yeast cultures was performed using an EVOS M5000 microscope (Thermo Fisher Scientific) at room temperature. Yeast cells were grown to the desired optical density under the specified growth conditions. Before imaging, cells were pelleted (4,000 × g for 5 min at room temperature), washed, and resuspended in sterile water. Subsequently, 3 µL of the dense yeast suspension was placed onto a glass slide for imaging.

### LD staining

For LD staining, cells were grown as described above, pelleted by centrifugation (4000 x g, 5 min), supernatant aspirated, and incubated for 5 min with either BODIPY 493/503 (Fisher Scientific, 11540326) or monodansylpentane (MDH; Abcepta, #SM1000b). After incubation, cells were washed once with water, pelleted again, and imaged. For vacuolar staining, cells were incubated for at least 30 min with the styryl dye FM 4-64 (Invitrogen, T13320) at a final concentration of 1 μM directly in the culture medium before collection for microscopic analysis.

### Quantitative image analysis

Images were analyzed using Fiji (ImageJ). Lipid droplets were quantified by counting per cell using the Multi-Point Tool. For DAG sensor localization, cells showing DAG signal at the ER or vacuole were manually counted. Identical acquisition and processing parameters were applied across samples.

### Lipolysis and lipophagy assays

For lipolysis, cells were precultured and subsequently diluted to 0.3 OD_600_ in SCD and grown for 48h to allow accumulation of TAG. At the 48h time point (T_0_), aliquots were harvested for baseline TAG quantification. These samples were immediately frozen in liquid nitrogen and stored at −80 °C prior to lipid extraction and analysis of TAG levels. To induce lipolysis, parallel 48h cultures were treated with cerulenin at a final concentration of 10 µg/mL and incubated for an additional 6h under standard growth conditions (30 °C, 225 rpm). After treatment (T_6_), cells were harvested and processed identically to T_0_ samples.

For lipophagy western blot assays, protein lysates from cell culture at indicated time points were prepared using a trichloroacetic acid (TCA) precipitation method. Briefly, aliquots corresponding to 10 OD_600_ units (where 1 ml of culture at 1.0 OD_600_ equals 1 unit) were collected in 15 mL culture tubes. Trichloroacetic acid was added to culture tubes at final concentration of 10% (w/v) and incubated for 20 min on ice or at −20 °C to allow protein precipitation. Precipitated proteins were collected by centrifugation at 15,000 × g for 3 min, transferred to 1.7 mL microcentrifuge tubes, and washed twice with 1 mL of ice-cold acetone. Washed pellets were air-dried in a non-heated speed vacuum for 10 min and resuspended in 200 μL of MURB buffer (50 mM sodium phosphate, 25 mM MES, pH 7.0, 1% SDS, 3 M urea, 0.5% 2-mercaptoethanol, and 1 mM sodium azide).

Cell disruption was performed by bead beating using acid-washed glass beads (approximately one-half pellet volume) for four cycles of 1 min beating followed by 30 s on ice at 4 °C. Lysates were incubated at 70 °C for 10 min and clarified by centrifugation at 10,000 × g for 1 min to remove unlysed cells and debris. A 10-μL aliquot of the resulting supernatant was used for SDS–PAGE and subsequent immunoblotting.

### Lipid extraction

Lipids were extracted using a modified Folch method (Folch et al., 1951) optimized for neutral lipids. Cell pellets were weighed, resuspended in sterile, deionized water, and disrupted with acid-washed glass beads using a mini bead beater (4 × 2 min, maximum speed, 4°C). Chloroform:methanol (2:1, v/v) was added, vortexed (5 minutes at room temperature), and transferred to a glass tube (Group A). The original tube was rinsed with methanol before combining with Group A. Additional chloroform was added, vortexed (1 minute), and centrifuged (2000 × g, 5 minutes, room temperature). The resulting lower phase was transferred to a new tube (Group B). Group A was re-extracted with 2:1 chloroform:methanol and the lower phase was combined with Group B. Sterile, deionized water was added to assist in phase separation as needed. Group B was washed twice with 1 M potassium chloride, vortexed, and centrifuged (2000 × g, 5 min, room temperature), discarding upper phases. The final organic phase was collected, dried under argon gas, and resuspended in chloroform (1 mg wet weight/µL). Samples (4–6 µL) were used for TLC analysis.

### Thin Layer Chromatography (TLC)

Silica gel plates were prebaked (145°C, 30 min), then marked with 18 lanes (1.5 cm outer margins, 1 cm between lanes) using a pencil. The development chamber was equilibrated with the appropriate solvent system: phospholipids: chloroform:acetone:methanol:acetic acid:water (50:20:10:15:5); neutral lipids: hexane: diethyl ether: acetic acid (80:20:10). Samples were applied and plates were developed at room temperature until lipid bands were fully separated. After air drying for 20 min, plates were sprayed with 3% copper (II) acetate in 8% phosphoric acid and charred at 145 °C for 1-8 hours to visualize bands; spraying and heating were repeated as needed.

### TLC quantification

Stained TLC plates were scanned and then processed for quantification using Fiji (ImageJ). Each plate contained a serial dilution of a standardized neutral lipid mixture. The standard mixture was prepared in chloroform to a final concentration of 10 mg/mL. The standardized neutral lipid mixture was used to create a standard curve in which the x axis displayed the calculated lipid mass in micrograms, and the y-axis displayed the estimated band intensity. Statistical analysis was performed in GraphPad Prism including ANOVA (*P > 0.05 (ns), < 0.05 (*), < 0.01 (**), < 0.001 (***), < 0.0001 (****)*).

### Western blot analysis

For SDS–PAGE analysis, 10 μL of cleared lysate was loaded per lane in 10% polyacrylamide gels and run at 115V for 40-60 min. Proteins were semi-dry transferred to a PVDF membrane before blocking in 5% non-fat milk in TBST (20 mM Tris-HCl pH 7.5, 150 mM NaCl, 0.1% Tween-20). Membranes were probed with αGFP primary antibody diluted in blocking solution (1:1000; Millipore-Sigma #11814460001). Detection was performed with HRP-conjugated secondary antibodies and ECL (BioRad Clarity™ Western ECL, #1705061) reagents, and signal was imaged on an iBright FL1500 system (Thermo Fisher). Total protein staining (No-Stain™ Protein Labelling Reagent, Fisher Scientific # A44449) served as the loading control.

## Acknowledgements

We thank Dr. Amra Saric and Dr. Juan Bonifacio for helpful discussions. We thank Dr. Cunqi Ye for critical reading of the manuscript. We thank Amit Joshi for sharing yeast tools. We thank Miriam Greenberg laboratory members for sharing supplies. H. Hariri is supported by the National Institutes of Health National Institute of General Medical Sciences (R35GM150892), Wayne State University start-up funds and University Research Award, Karmanos Cancer Institute – Strategic Research Initiative Grant (KCI-SRIG), Richard Barber Interdisciplinary Research Program. The Microscopy, Imaging and Cytometry Resources Core is supported in part by NIH Karmanos Cancer Institute Center Grant (P30 CA22453) and Wayne State University.

The authors declare no competing financial interest.

## Author contributions

H. Hariri conception and design of the study, data interpretation, drafting and revising the manuscript. D. Adebayo, lipid analysis and qPCR. E. Obaseki, lipophagy and lipolysis assays. S. Miller, E. Obaseki, Z. Saadeh, imaging and image quantification. D. Adebayo, E. Obaseki, S. Miller, figure preparation, critical reading of the manuscript. M. Aboumourad, Z. Pomicter, L. Kreinbring, M. Rahal, K Vasudeva, G. Frost, R. Smadi, generating yeast strains, technical and editorial assistance.

## Supplementary Figures

**Figure S1.** Nvj3-LD colocalization is independent of Mdm1 and its deletion increases LD numbers under nutrient stress. a) Fluorescent imaging of overexpressed Nvj3–GFP (green) and LDs (magenta, MDH) in *mdm1Δ* cells grown in SCD + 0.2% oleate (OA). Nvj3 remains enriched at LDs in the absence of Mdm1. Scale bar, 1 µm. b) Fluorescence micrographs showing Erg6-GFP-labeled LDs (green) in wild-type (WT) and *nvj3Δ* cells at log and after 6h of acute glucose starvation (AGS; 0.001 % glucose). Corresponding transmitted-light (DIC) images are shown. Scale bars, 3 µm. c) Quantification of Erg6-GFP puncta per cell in WT and *nvj3Δ* yeast in (b) at the indicated growth conditions (log and AGS). Data represent mean ± SD; *n* ≥ 100 cells. *P* < 0.05 (*), ns = not significant (Student’s *t* test).

**Figure S2.** Neutral lipid and phospholipid TLC and quantifications in WT and nvj3Δ. a) Thin-layer chromatography (TLC) analysis of neutral lipid profiles from wild-type (WT) and *nvj3Δ* cells under logarithmic (Log), stationary phase (SP; 72h), and acute glucose starvation (AGS; 0.001 % glucose, 6h). Bands corresponding to sterol esters (SE), triacylglycerols (TAG), free fatty acids (FFA), ergosterol (ERG), and diacylglycerols (DAG) are indicated. b) Quantification of ergosterols in (a). *nvj3Δ* cells exhibit increased accumulation of ERG relative to WT during AGS. c) TLC analysis of phospholipid species in WT and *nvj3Δ* cells under log and AGS conditions. Labeled bands correspond to phosphatidic acid (PA), phosphatidylethanolamine (PE), phosphatidylcholine (PC), and phosphatidylinositol (PI). d) Quantification of phospholipid species in (c) from wild-type (WT) and *nvj3Δ* cells grown in log measured by TLC. Mean ± SD from three biological replicates. *P* < 0.05 (\**), < 0.01 (*\*\**), < 0.001 (*\*\*\*), < 0.0001 (****); Two-way ANOVA.

**Figure S3.** Lipophagy and LD-vacuole colocalization not impaired in *nvj3Δ* cells. a) Quantification of LDs localized within vacuoles during log and stationary phase (SP) growth, categorized as 0, 1-3, or >3 LDs per vacuole. No significant differences were observed between WT and *nvj3Δ*. Mean ± SD; *n* ≥ 100 cells. b) Representative fluorescence micrographs quantified in (A) of LD (Erg6-GFP) and vacuole (FM4-64) colocalization in WT and *nvj3Δ* cells under log and SP. Scale bar, 3 µm. c) Western blot analysis of Erg6-GFP cleavage in WT and *nvj3Δ* during lipophagy induction in nitrogen starvation (SD-N). Free GFP bands released upon vacuolar degradation are indicated. Total protein staining is used as a loading control. d) Quantification of Erg6-GFP degradation from (c), comparing the extent of GFP cleavage between WT and *nvj3Δ.* Mean ± SD from three biological replicates.

**Figure S4.** Loss of Nvj3 alters phospholipid remodeling during induced TAG synthesis. A) Thin-layer chromatography (TLC) analysis of phospholipid species from *pGAL-DGA1* (ΔLD) and *pGAL-DGA1-nvj3Δ* (ΔLD-nvj3Δ) cells collected 2, 8, and 14h post-galactose induction. Bands corresponding to phosphatidic acid (PA), phosphatidylethanolamine (PE), phosphatidylcholine (PC), and phosphatidylinositol (PI) are indicated. B) Quantification of PA species in (a). *nvj3Δ* cells exhibit reduced PA C) Quantification of PC, PE, PI species in (a). PC and PE remain relatively unchanged, PI increases in ΔLD-nvj3Δ. D) TLC analysis of phospholipid species from *pGAL-LRO1* (ΔLD) and *pGAL-LRO1-nvj3Δ* (ΔLD-*nvj3*Δ) cells harvested 2, 8, and 14h post-galactose induction. Bands corresponding to PA, PC, PE, and PI are shown. E) Quantification of PA species in (d) where *nvj3Δ* cells display reduced PA. F) Quantification of PC, PE, PI species in (d). *nvj3Δ* cells display increase in PC and PI while PE is relatively unchanged. Mean ± SD, n = 2. *P < 0.05 (*), < 0.01 (**), < 0.001 (***)*; Two-way ANOVA.

**Figure S5.** Kinetics of LD formation in Dga1– and Lro1-driven systems in the absence of Mdm1 and Nvj3. a) Fluorescence micrographs of BODIPY staining 2h and 24h post-galactose induction in *pGAL-DGA1* (ΔLD and ΔLD-*nvj3*Δ). Scale bar, 3 µm. b) Quantitative changes in LD overtime in *pGAL-DGA1* (ΔLD and ΔLD-*nvj3*Δ) system. Percentage of cells with at least 1 LD at different timepoints. Mean ± SD, *n* ≥ 100 cells; *P < 0.05 (*), < 0.01 (**), < 0.0001 (****),* ns = not significant; Two-way ANOVA. c) Fluorescence micrographs of BODIPY staining 2h and 8h post-galactose induction in *pGAL-LRO1* (ΔLD and ΔLD-*nvj3*Δ). Scale bar, 3 µm. d) Quantitative changes in LD overtime in *pGAL-LRO1* (ΔLD and ΔLD-*nvj3*Δ) system. Percentage of cells with at least 1 LD at different timepoints. Mean ± SD, *n* ≥ 100 cells; *P < 0.05 (*), < 0.01 (**), < 0.0001 (****),* ns = not significant; Two-way ANOVA. e) Fluorescence micrographs of BODIPY staining at 4 and 6h post-galactose induction in *pGAL-DGA1* (ΔLD), *pGAL-DGA1-mdm1Δ* (ΔLD-*mdm1*Δ), and *pGAL-DGA1-nvj3Δ* (ΔLD-*nvj3*Δ). Scale bar, 3 µm. f) Quantification of images (e). Number of LDs per cell at 6h (Left). Percentage of cells with at least 1LD at 6h (Left). Mean ± SD, *n* ≥ 100 cells; *P < 0.05 (*), < 0.01 (**), < 0.0001 (****),* ns = not significant; Two-way ANOVA.

**Figure S6.** Dga1 contributes to LD accumulation in *nvj3Δ* cells and DAG mislocalization persists under oleate loading. a) Fluorescent imaging of GFP-tagged DAG sensor (C1-domain from Protein Kinase D) in wild-type and *nvj3*Δ cells grown in oleate-supplemented (SCD 0.01% + OA) conditions. Scale bar, 3 µm. b) Quantification of (a) DAG localization. (Top) Percent cells with vacuolar DAG signal. (Bottom) Percent cells with nER (nuclear ER) DAG signal; in wildtype (WT and *nvj3*Δ cells. *n* ≥ 100 cells. c) Relative mRNA expression of *LRO1* in wild-type and *nvj3*Δ cells during log phase and acute starvation. Mean ± SEM, n = 4; *P < 0.05 (*), < 0.01 (**);* Two-way ANOVA. d) Fluorescent imaging of Dga1-GFP overexpression via low-copy plasmid in wild-type and *nvj3*Δ cells grown in rich (SCD 2%) and low-carbon (SCD 0.001%) conditions. Scale bar, 3 µm. e) Fluorescence micrographs showing AUTODOT-stained LDs in wild-type, *nvj3Δ*, *nvj3Δ lro1Δ*, and *nvj3Δ dga1Δ* cells during log phase and after acute glucose starvation (AGS). Scale bar, 3 µm. f) Quantification of total LD fluorescence intensity per cell in wild-type, *nvj3*Δ, *nvj3*Δ *lro1*Δ, and nvj3Δ dga1Δ strains during log and starvation phases imaged in (A). Mean ± SD, *n* ≥ 100 cells; *P < 0.05 (*), < 0.001 (***), < 0.0001 (****),* ns = not significant, Two-way ANOVA.

## Supplemental Tables

**Table S1:**
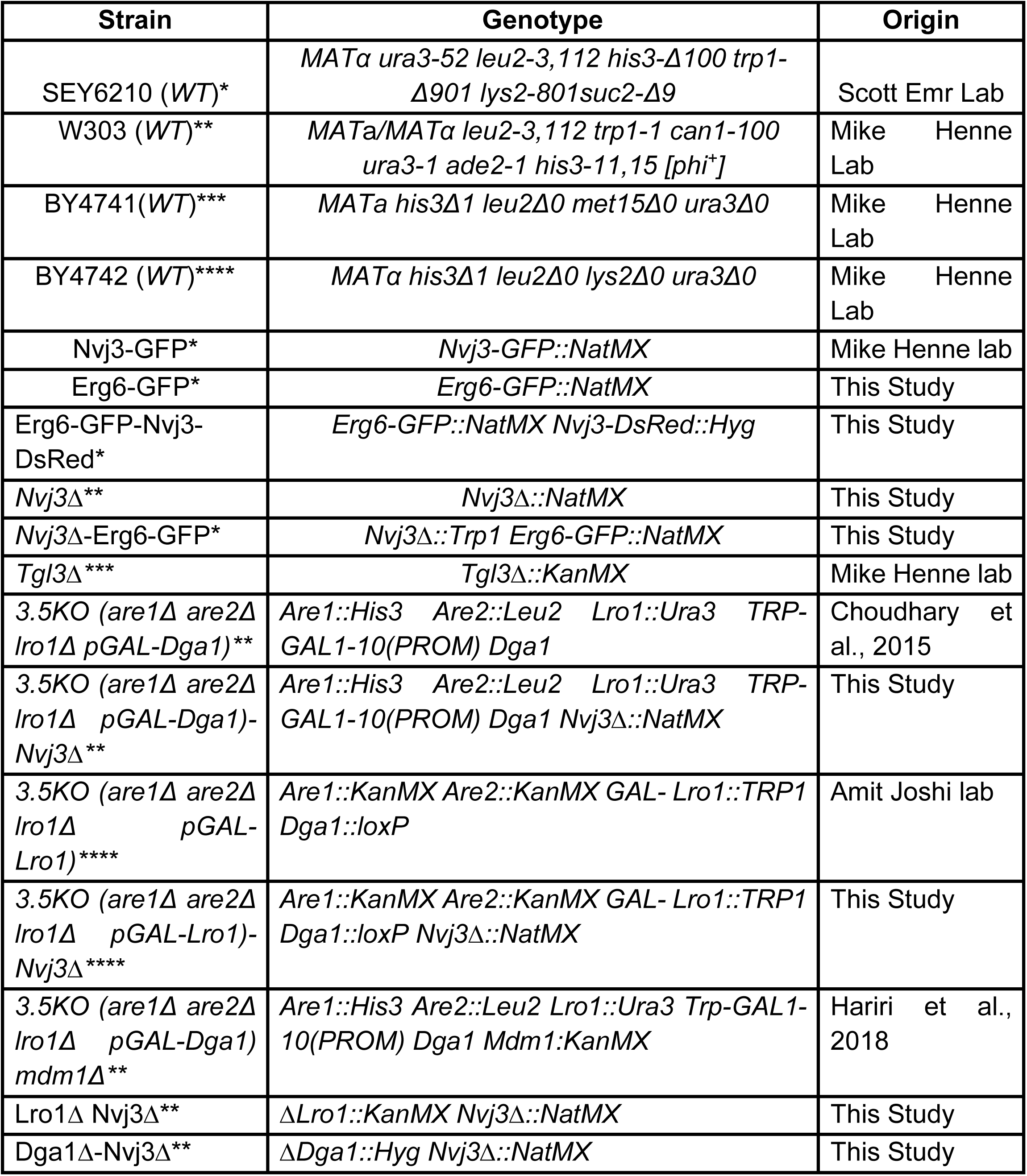
List of yeast strains used in this study.

**Table S2:** Yeast expression plasmids used in this study.

| Plasmid | Characteristics | Origin |
| --- | --- | --- |
| pBP73G (EV) | Empty pBP73G (*URA3*, GPD promoter) | Mike Henne Lab |
| Nvj3-GFP (pBP73G) | Nvj3-GFP from pBP73G (*URA3*, GPD promoter) | Mike Henne Lab |
| Ycp111-Dga1-GFP | Centromeric *LEU2* plasmid encoding Dga1-GFP; AmpR bacterial selection marker | Amit Joshi Lab |
| Yep181-Dga1-GFP | High-copy *LEU2* plasmid encoding Dga1-GFP; AmpR bacterial selection marker | Amit Joshi Lab |
| GFP-C1 DAG sensor | LEU2-marked yeast expression plasmid. DAG-binding tandem C1 domains of protein kinase D (C1a/b-PKD) fused to GFP and the transmembrane domain of Ubc6, a tail-anchored ER protein. | Choudhary et al., 2018 |

**Table S3:**
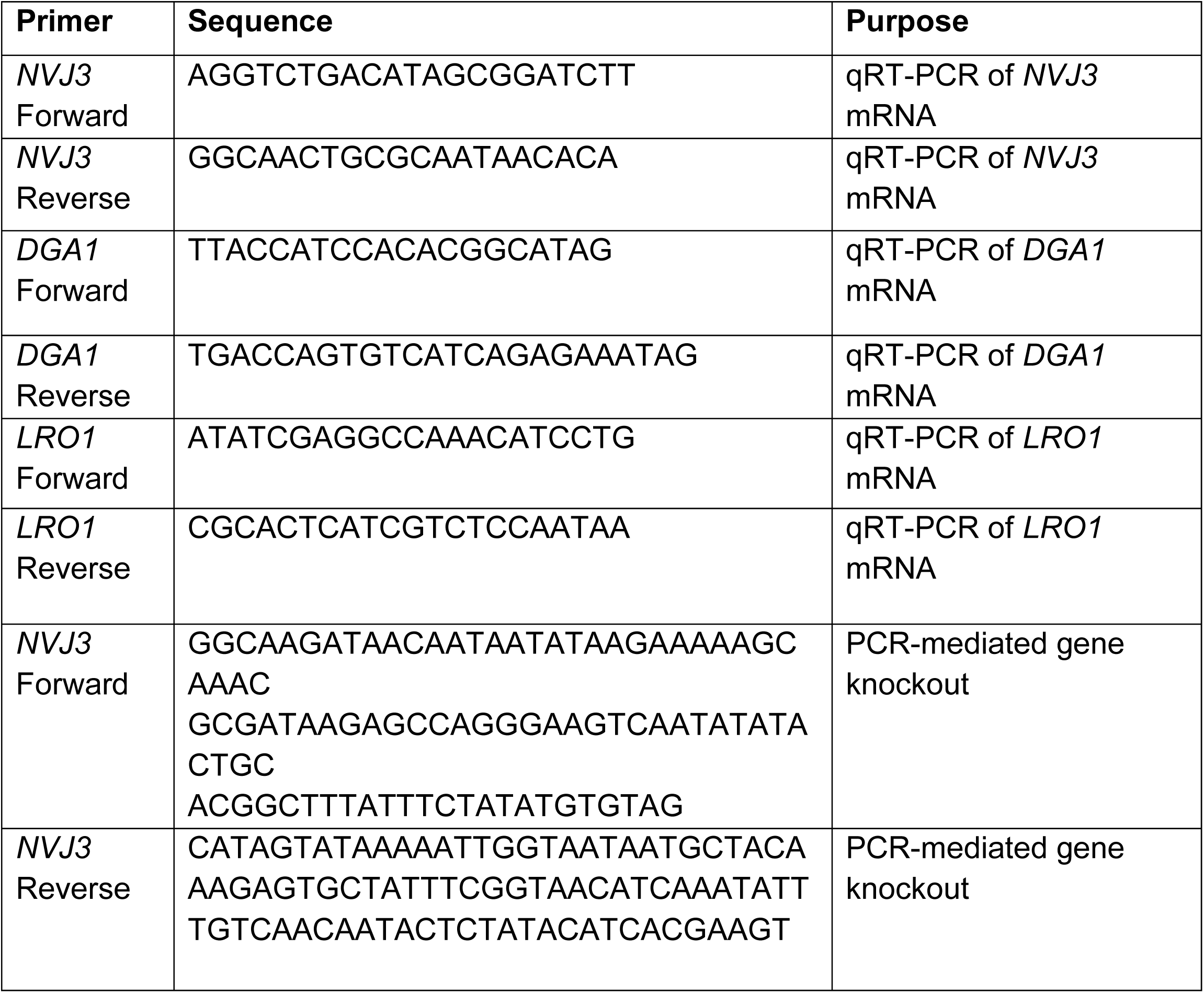
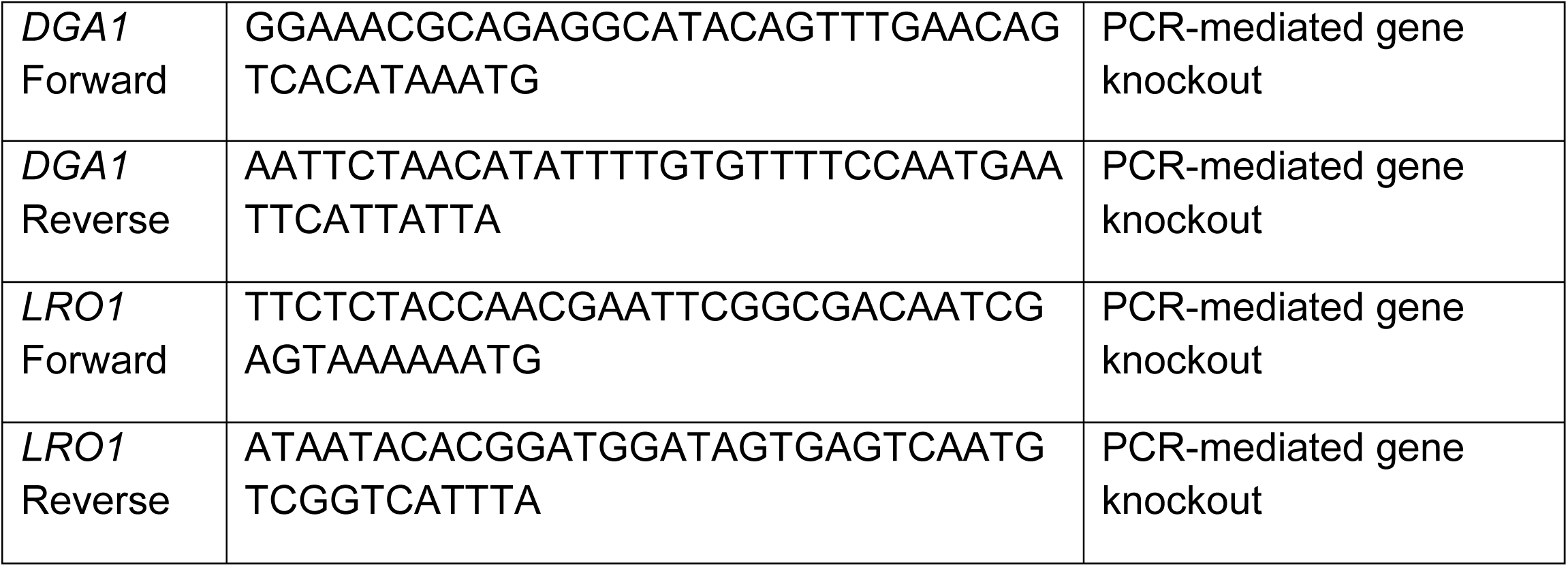
List of primers used in this study.

