## Supplementary figures and images for "Nvj3 regulates Dga1-mediated triacylglycerol synthesis and lipid droplet formation at ER contact sites"

### Supplemental figures

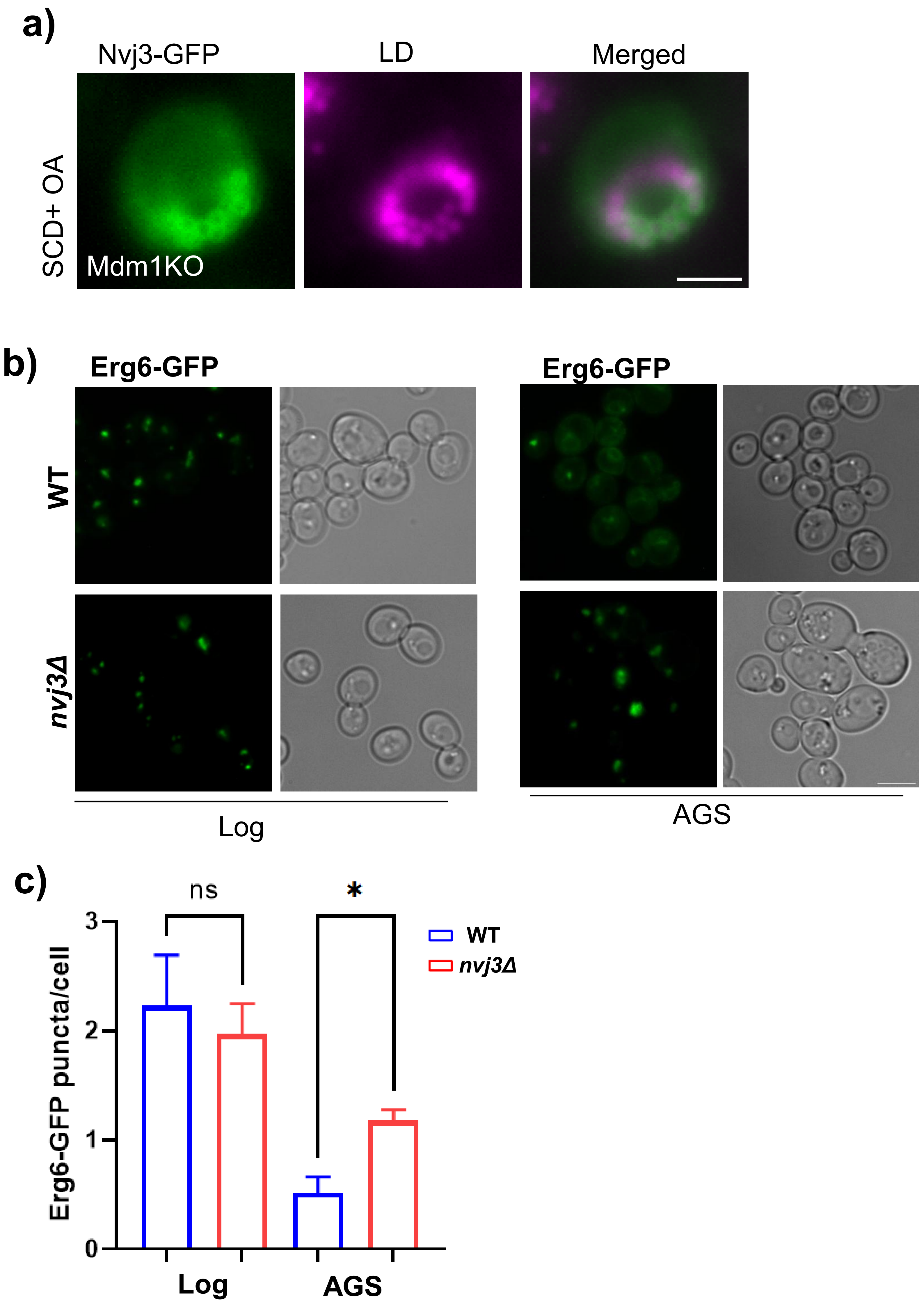

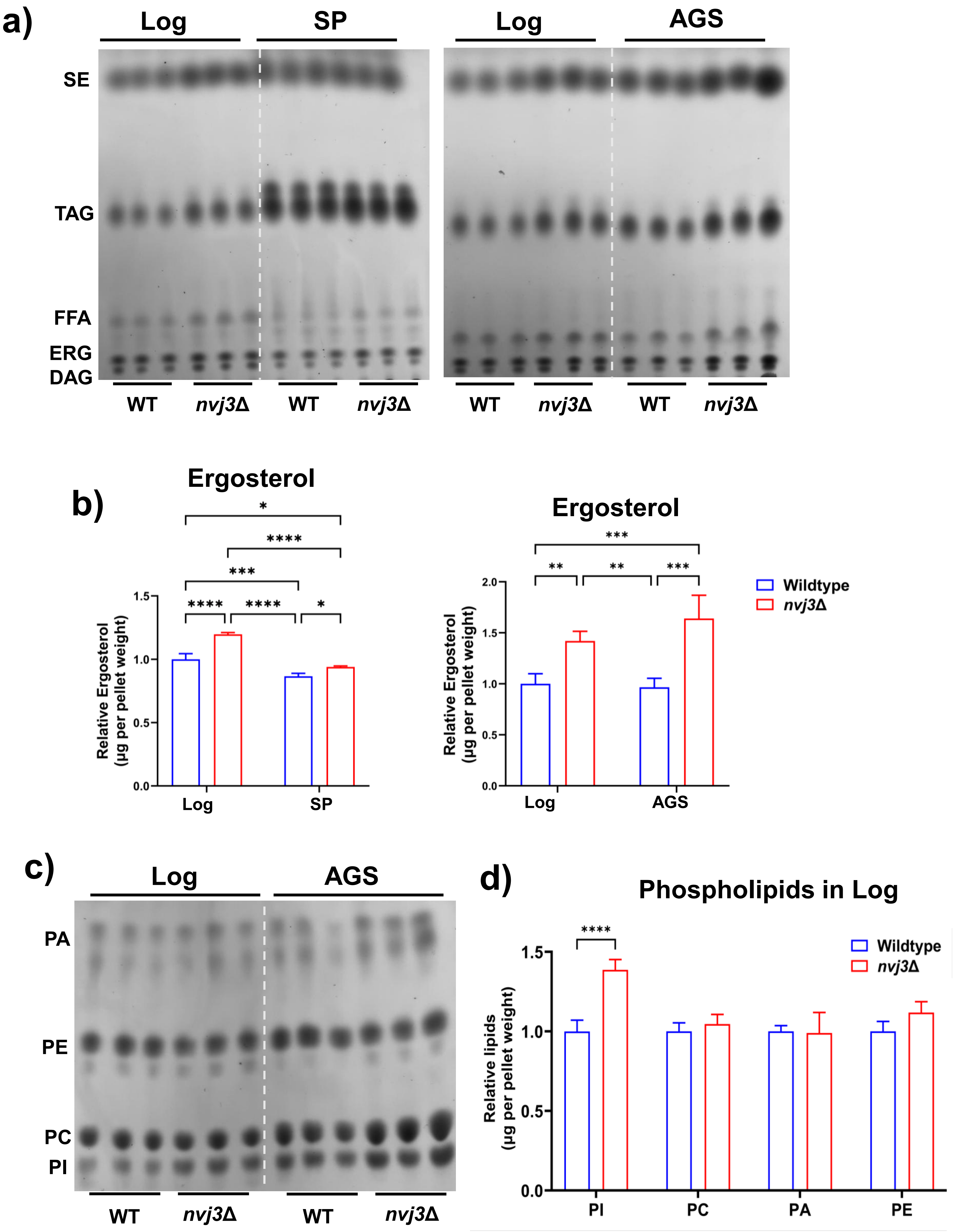

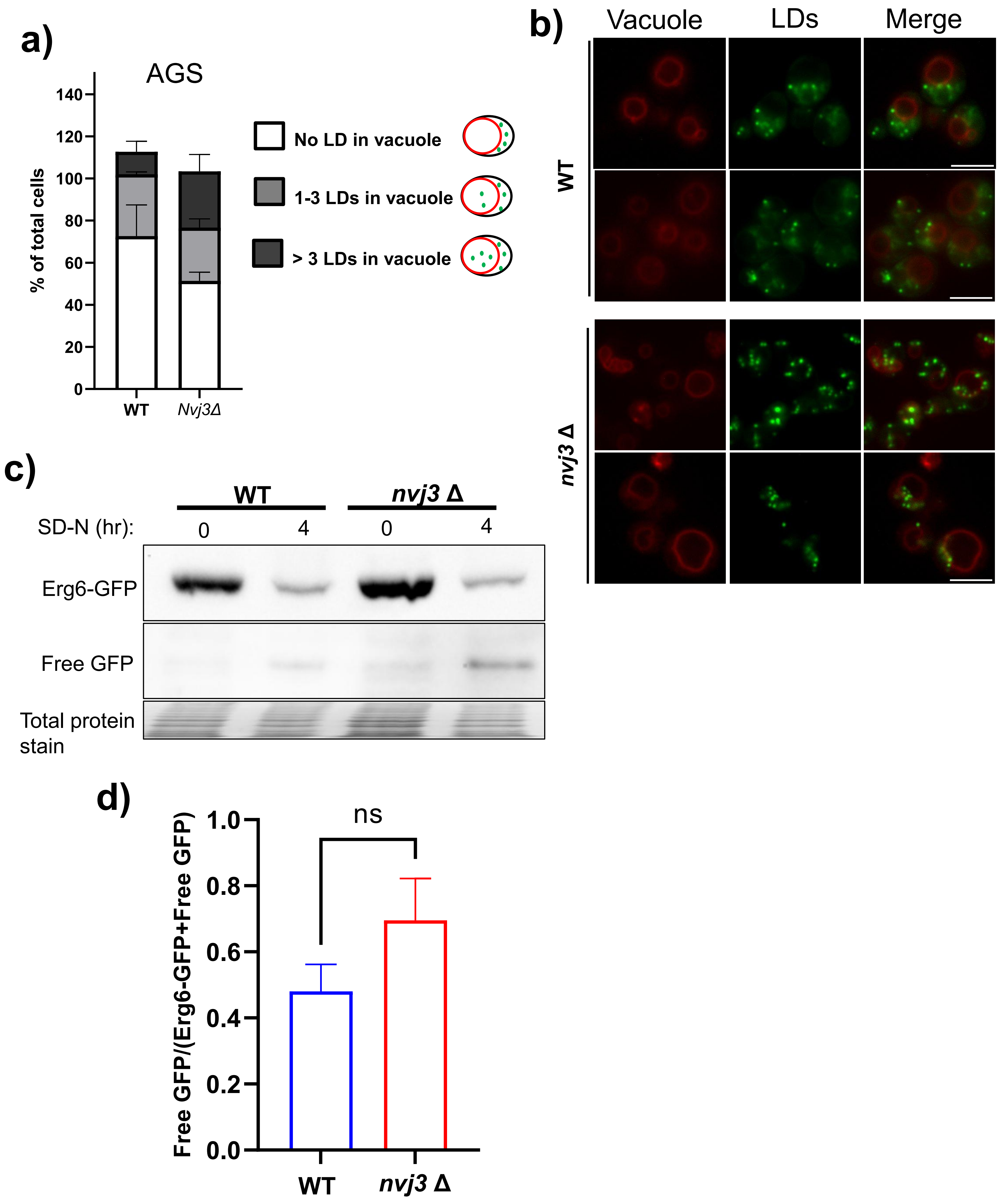

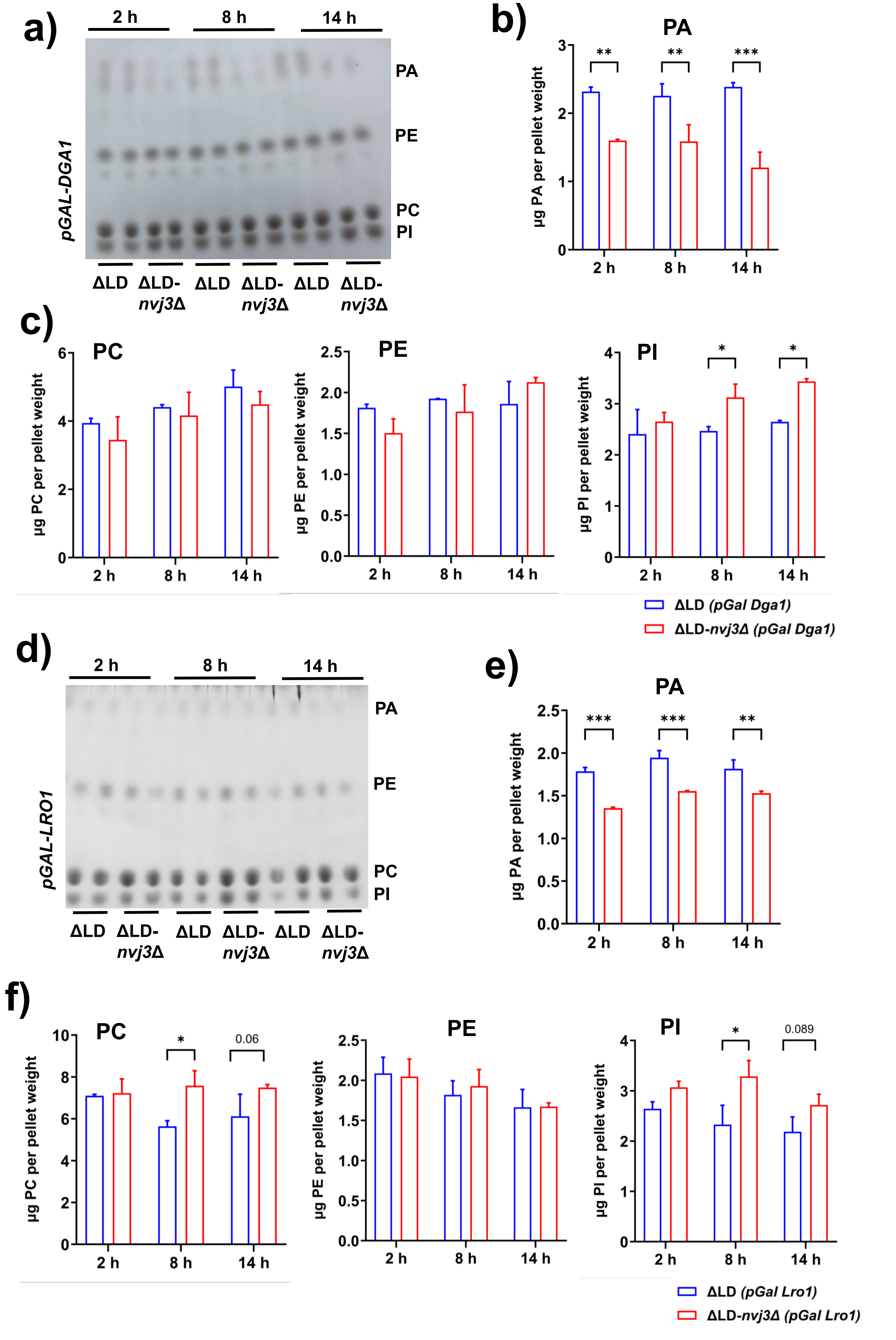

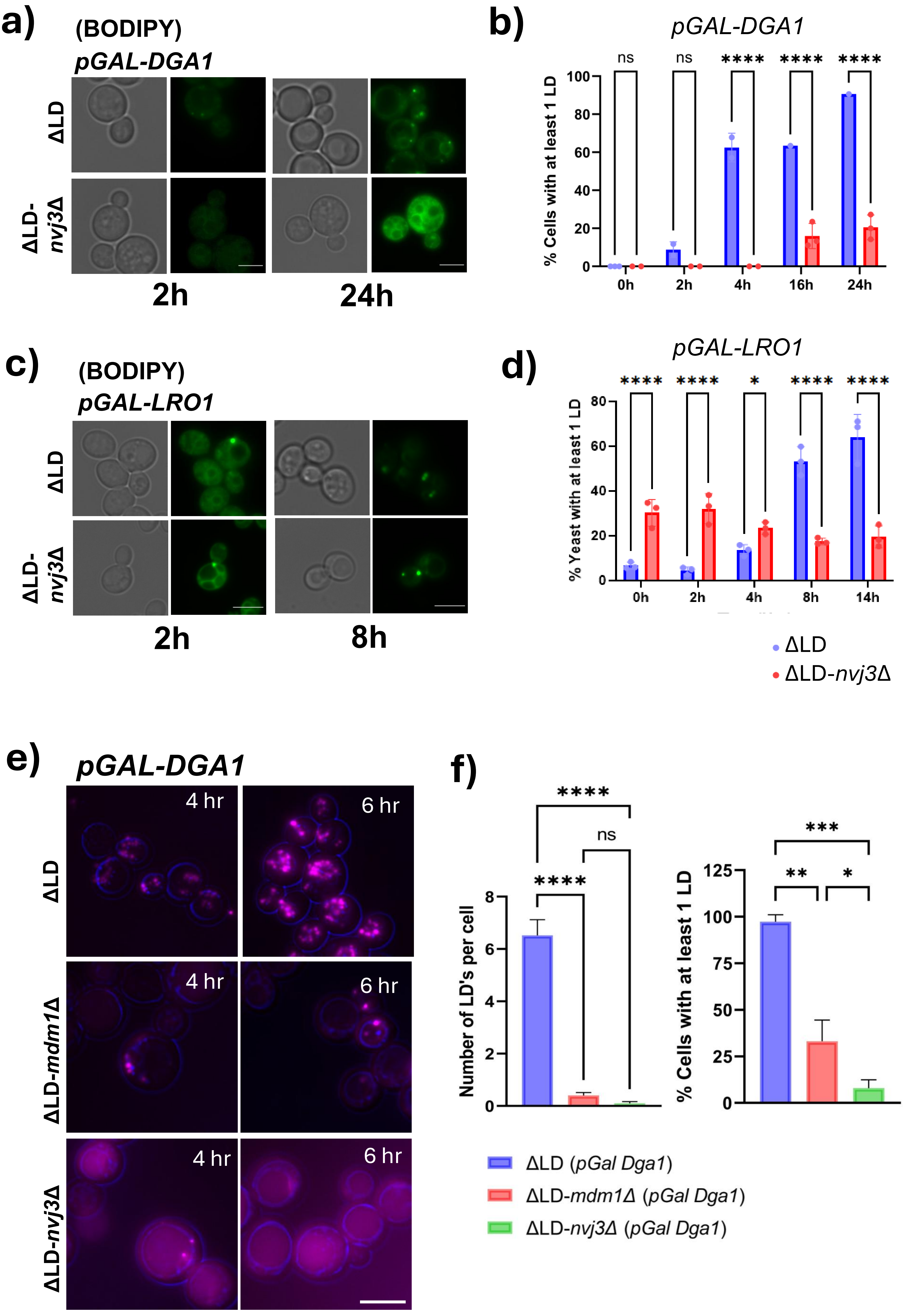

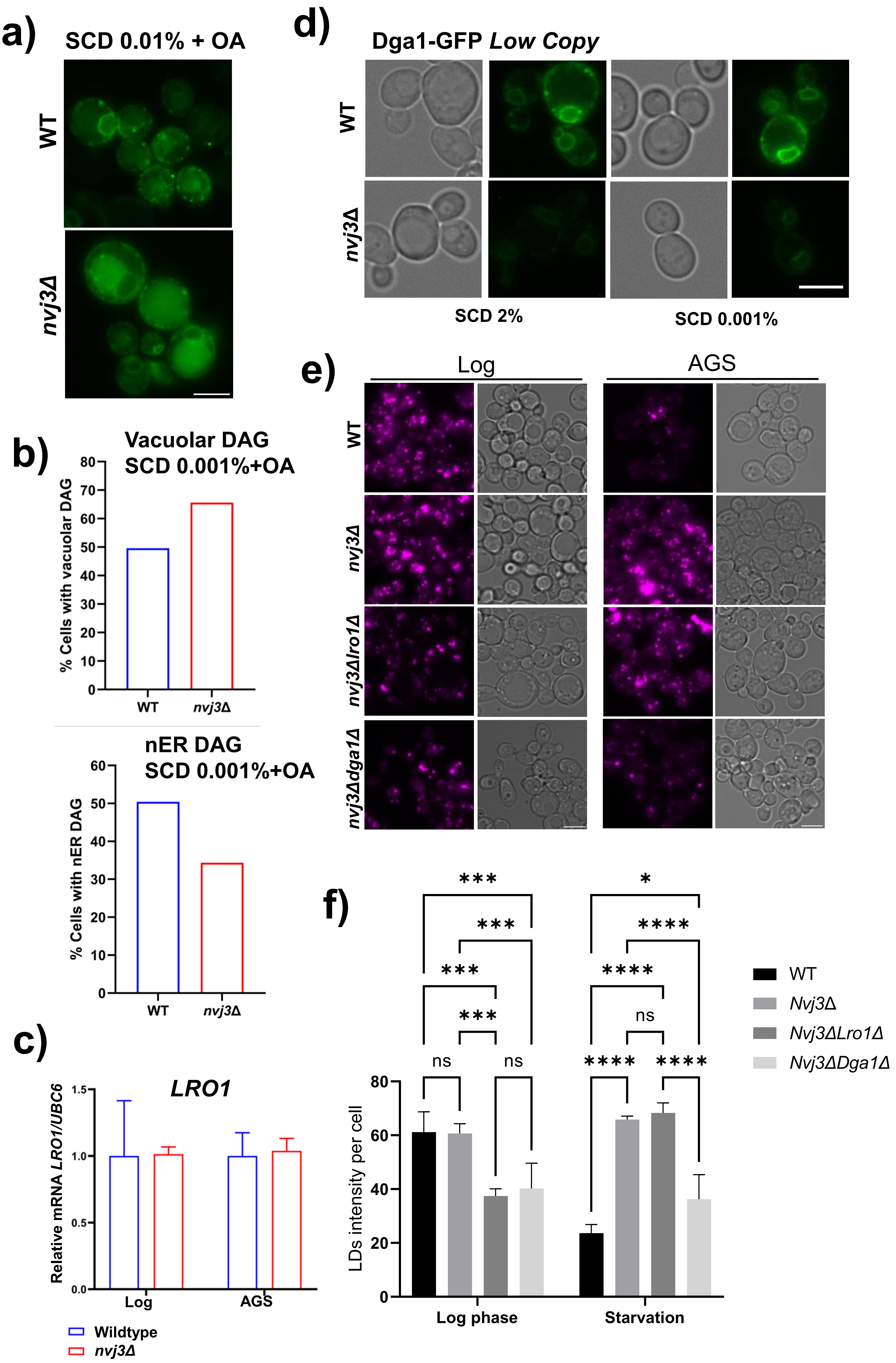
